# Endothelial cell stiffness and type drive the formation of biomechanically-induced transcellular pores

**DOI:** 10.1101/2024.10.23.619950

**Authors:** Seyed Mohammad Siadat, Haiyan Li, Brigid A. Millette, Babak N. Safa, Cydney A. Wong, M. Reza Bahrani Fard, Sietse T Braakman, Ian Tay, Jacques A. Bertrand, A. Thomas Read, Lisa A. Schildmeyer, Kristin M. Perkumas, W. Daniel Stamer, Darryl R. Overby, C. Ross Ethier

## Abstract

Formation of transcellular pores facilitates the transport of materials across endothelial barriers. In Schlemm’s canal (SC) endothelium, impaired pore formation is associated with glaucoma. However, our understanding of the cellular processes responsible for pore formation is limited by lack of *in vitro* assays. Here, we present a novel platform for studying transcellular pore formation in human endothelial cells. We induced pores in SC cells by seeding them atop micron-sized magnetic beads followed by application of a magnetic field to subject cells to a basal to apical force, mimicking *in vivo* biomechanical forces. The pore formation process was dynamic, with pores opening and closing. Glaucomatous cells exhibited impaired pore formation that correlated with their increased stiffness. We further discovered that application of forces from the apical to basal direction did not induce pores in SC cells but resulted in formation of pores in other types of endothelial cells. Our studies reveal the central role of cell mechanics in formation of transcellular pores in endothelial cells, and provide a new approach for investigating their associated underlying mechanism/s.

## Main

Transcellular transfer of materials (*e.g.*, cells, proteins, or fluids) directly across cellular barriers occurs during leukocyte transmigration through microvascular endothelial cells,^1–7^ glomerular filtration across endothelial cells,^8^ and drainage of aqueous humor (AH) from the eye via pores within the hybrid blood vasculature/lymphatic endothelial cells of Schlemm’s canal’s (SC) inner wall.^9–12^ In these cases, a pore (opening) must be created through an endothelial cell without damaging the cell. Our understanding of the cellular processes responsible for such pore formation is limited, in part, due to the small size of these pores and lack of suitable *in vitro* assays replicating the pore formation process.

One important application for such assays is to screen for novel therapies to treat glaucoma, the leading cause of irreversible blindness^13^ affecting over 70 million individuals globally.^14,15^ All existing medical and surgical treatments for glaucoma aim to lower intraocular pressure (IOP) to decelerate disease progression;^16,17^ yet, existing approaches are insufficient.^18^ IOP is strongly influenced by AH transport through micron-sized pores that form in endothelial cells lining the inner wall of SC, the only continuous cellular barrier within the major pathway for drainage of AH from the eye^19^ (**Fig. 1a**). Recent discoveries showing that SC pore formation is a mechanosensitive process and that pore formation can be facilitated by mechanical stretching^20^ and transendothelial perfusion,^21^ motivate the development of discovery tools to study the mechanism(s) of mechanosensitive pore formation. Two types of pores form in SC cells and participate in fluid drainage from the eye: transcellular and paracellular^12^ (**Fig. 1a**).

**Fig. 1.**
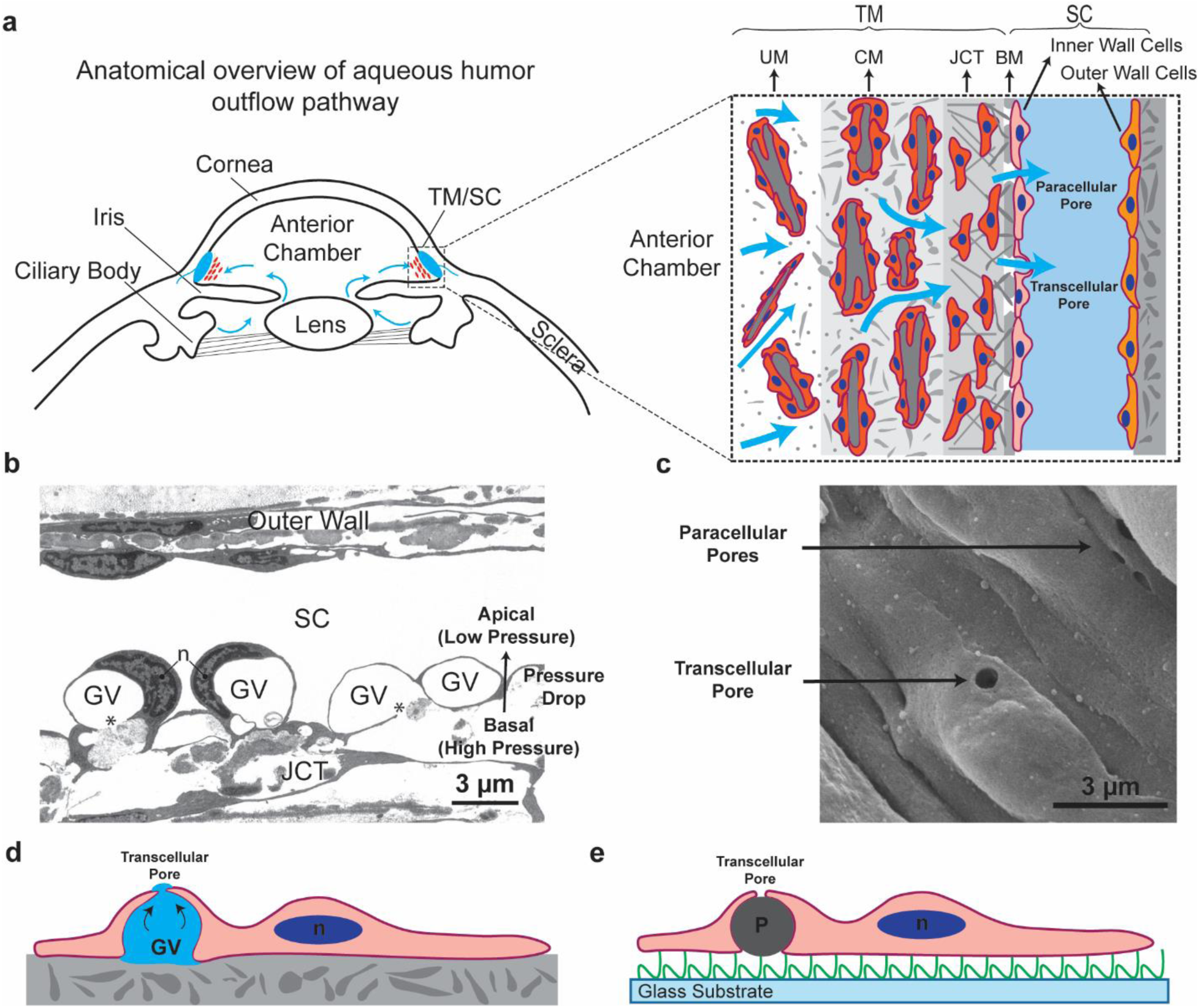
Pore formation in SC inner wall cells. **a,** Anatomical overview of aqueous humor (AH) outflow pathway. AH is produced by the ciliary body, flows (shown by blue arrows) into the anterior chamber, and exits the chamber primarily through the trabecular meshwork (TM) and Schlemm’s canal (SC). The magnified inset shows AH flow through consecutive regions of TM: uveal meshwork (UM), corneoscleral meshwork (CM), and juxtacanalicular tissue (JCT). AH then crosses SC inner wall cells through transcellular or paracellular pores. Basement membrane (BM) and extracellular matrix components are shown in grey. **b,** A transmission electron micrograph showing significant deformation experienced by SC inner wall endothelial cells during giant vacuole (GV) formation. GVs are outpouchings of the endothelial cells lining SC, protruding into the canal’s lumen and creating a fluid-filled space between the cell and the underlying BM (adopted with permission from Pedrigi *et al*.^27^). ‘n’: Nucleus; ‘*’: Opening in BM. **c,** Scanning electron micrograph of the SC inner wall as observed from inside the canal. GVs are often associated with development of transcellular pores,^28^ which create transcellular routes for drainage of AH from anterior chamber of the eye into the SC lumen. Note the bulging structures (GVs and nuclei) and pores (arrow). SC from human donor eyes (53-year-old female). **d,** Schematic illustrating transcellular pore formation in a SC cell *in vivo* exposed to significant stretch due to the pressure drop across the inner wall, during GV formation. **e,** Schematic illustrating transcellular pore formation in a SC cell induced by a particle *in vitro*. In our *in vitro* setup, phSC cells are seeded over a gelatin coated glass substrate and ~ Ø 5 µm carboxyl-coated, ferromagnetic particles to create local cellular deformation. The particles are about the same size as the GVs *in vivo*. Our hypothesis posits that as cells spread over these particles, they become thinner which ultimately gives rise to transcellular pores. ‘P’: particle; green: gelatin substrate underlying cells in culture; blue: cell nuclei.

Transcellular pores are triggered by the significant deformation experienced by SC endothelial cells in response to the normal transcellular basal-to-apical pressure drop across the monolayer^22^ *in vivo* (**Fig. 1b and c**). Further, SC pores are reduced in glaucoma,^23,24^ motivating the discovery of “SC-active agents” for IOP control. We here describe an innovative *in vitro* platform enabling generation and detection of transcellular pores in cultured primary human SC cells. Using this platform we have conducted basic science studies showing that pore formation is tightly associated with cell stiffness. This platform offers the opportunity for further studies on the mechanobiology of pore formation, on molecular participants in the pore forming process, and, eventually, for high throughput screening of compounds that promote pore formation.

## Results

### Local basal-to-apical cellular stretch induces transcellular pore formation in SC cells

Primary human SC (phSC) cells were isolated from *post mortem* donor eyes, cultured and characterized as previously described.^25,26^ We postulated that seeding phSC cells on top of micron-sized particles would induce localized cellular deformation, mimicking the *in vivo* basal-to-apical biomechanical loading (**Fig. 1d and e**). This, in turn, was expected to initiate the formation of transcellular pores. We thus randomly scattered carboxyl-coated, ferromagnetic microspheres with nominal Ø 5 µm on gelatin-coated glass substrates, followed by seeding phSC cells on top of these particles (see Methods). Cells were then cultured for 2 to 7 hours so that they adopted a spread configuration similar to that seen *in situ*.

Scanning electron micrographs of fixed cells confirmed that phSC cells stretched over particles (**Fig. 2a**), with transcellular pores appearing above some particles (**Fig. 2b**). We denote pores that formed in association with particles as “particle-induced pores.” Notably, these transcellular pores exhibited smooth edges, allowing for clear differentiation from artifacts characterized by irregular and jagged edges that were likely byproducts of the scanning electron microscopy (SEM) sample preparation process.^29^ Using high resolution SEM images, cell membrane thickness at the smooth edge of pores were measured as 56 ± 11 nm (mean ± standard deviation, N = 15 cells). Note that the transcellular pores were also formed at the location of smaller (Ø 2 µm) particles (**Fig. S1**); however, we only used Ø 5 µm particles in the rest of this study to facilitate detection of particles, and hence of pores overlying particles.

**Fig. 2.**
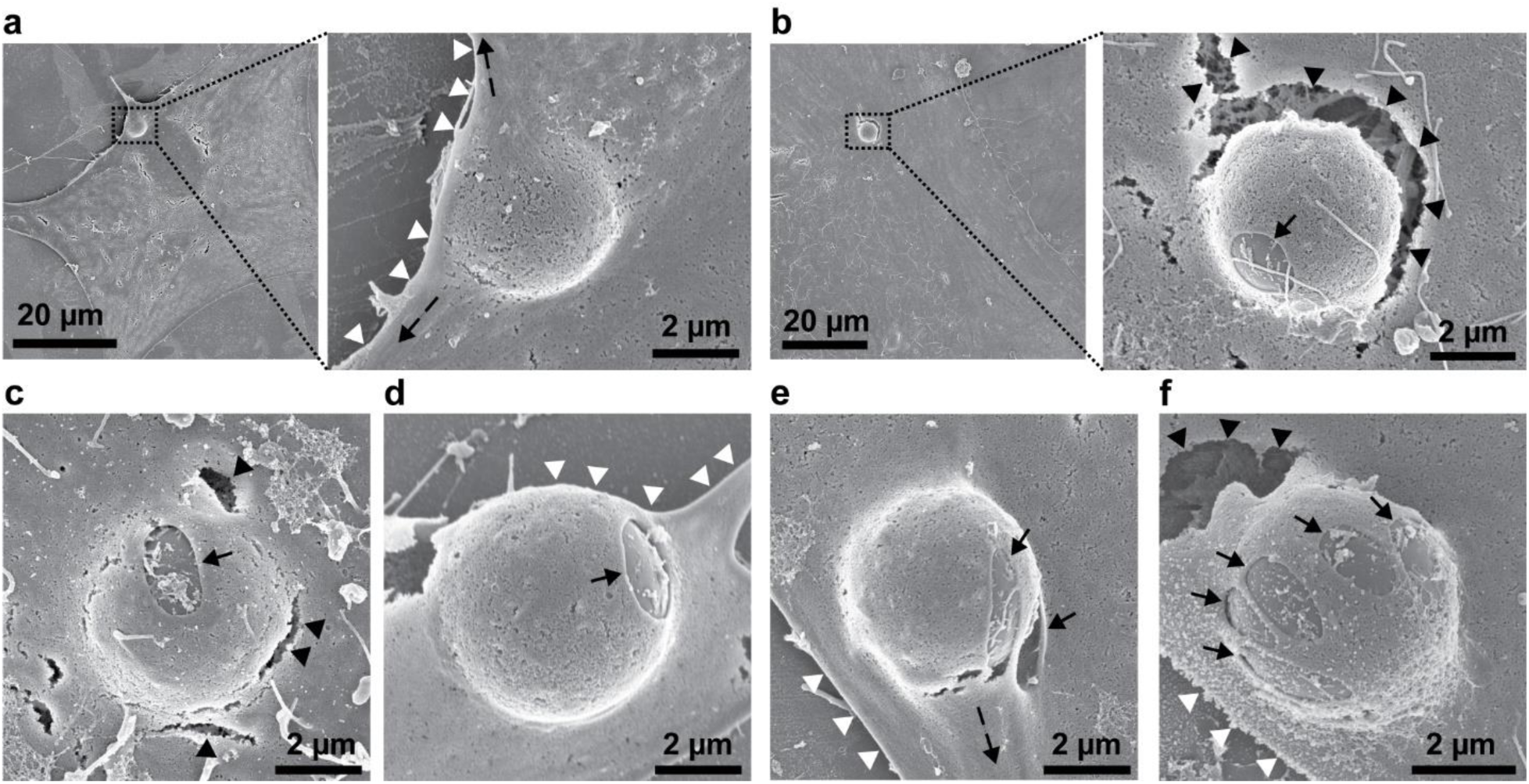
Scanning electron micrographs demonstrating particle-induced cell deformation and transcellular pore formation. **a** and **b,** Left panels show low-magnification images of phSC cells overlying particles, while right panels show higher-magnification views of the insets (black boxes), highlighting cell deformation at the particle locations. **c-f,** SEM micrographs demonstrating various aspects of particle-induced transcellular pores. In panel **a**, we observe stretching of a phSC cell (suggested by black dashed arrows) associated with a particle near the cell’s border (white arrowheads). Although this particle did not induce the formation of a transcellular pore, noticeable cellular deformation around the particle is evident. Panel **b** shows a transcellular pore (black arrow) associated with a particle. Note the artifactual cracks (black arrowheads) with irregular, jagged edges near the particle. **c,** A transcellular pore formed directly above a particle, with several noticeable cracks in proximity. **d,** A transcellular pore formed to the side of the particle, situated near the cell border. **e,** Two transcellular pores formed on the side of a particle, elongated in the same direction as the cell stretched. **f,** Multiple transcellular pores formed above a particle, resulting in a “toffee-like” structure. Black arrows indicate transcellular pores, black arrowheads indicate artifactual pores or cracks with jagged edges, white arrowheads indicate cell borders, and black dashed arrows indicate the apparent direction of cell stretching.

We observed a significant diversity of transcellular pore locations and morphologies. For example, some pores formed directly above (**Fig. 2c**) or adjacent to (**Fig. 2d**) particles. Further, it appeared that particles could induce pore formation irrespective of their specific location beneath cells; for example, transcellular pores formed in cells both with particles situated away from (**Fig. 2b and c**) and near (**Fig. 2d and e**) the cell border. In many cases, we observed distinct and obvious artifactual cracks, particularly adjacent to the particles (**Fig. 2b and c**), which we speculate is due to the cells stretching over the particles, rendering them more vulnerable to artifacts during the SEM sample preparation process.^29^ However, there were instances (**Fig. 2e**) where the cell did not exhibit major artifacts adjacent to a particle despite noticeable cellular deformation. In the case shown in **Fig. 2e**, transcellular pores were formed to the side of the particle and extended along the direction of membrane stretching. Further, some particles induced the formation of only one transcellular pore (**Fig. 2b, c, and d**), while others induced multiple transcellular pores (**Fig. 2e and f**). Of note, the transcellular pores shown in **Fig. 2f** exhibit a structure reminiscent of “toffee-like” pores previously reported by Braakman *et al*.^20^

### Transcellular pores are rapidly detectable using an *in vitro* fluorescent assay

We employed confocal microscopy and an established fluorescent assay^30–32^ (**Fig. 3**) to further investigate cell deformation at the location of particles and to also rapidly identify transcellular pores without the need for time-consuming SEM imaging. In this assay, a fluorescent tracer (fluorescently labeled streptavidin) adheres to a biotinylated gelatin substrate at sites not obscured by cells, marking areas surrounding subconfluent cells and, notably, at transcellular pore locations (**Fig. 3a, b**).

**Fig. 3.**
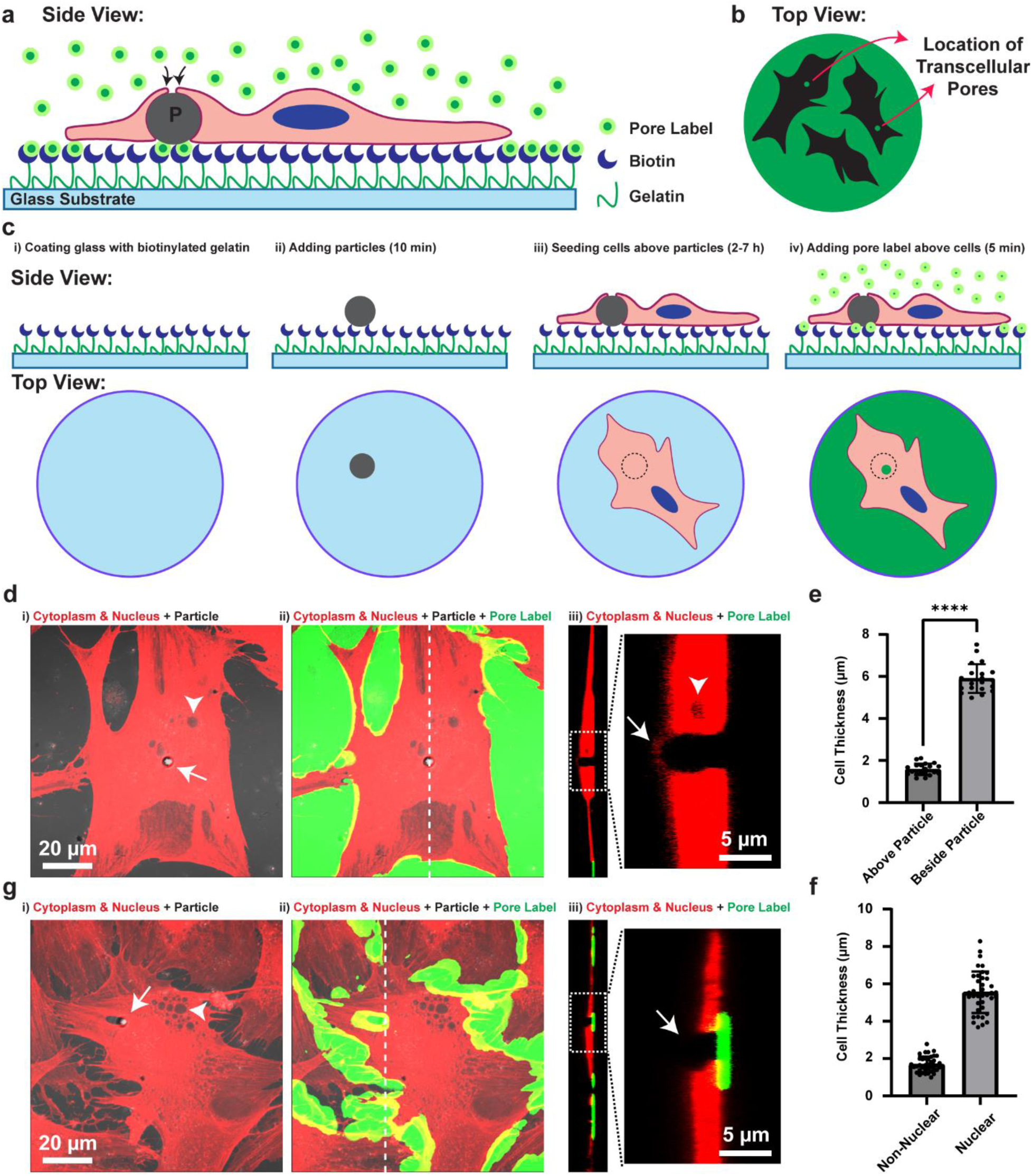
Cell deformation induced by particles and transcellular pore detection without SEM imaging. **a,** Schematic illustrating the fluorescent assay (side view) for rapid transcellular pore detection. phSC cells are grown on biotinylated gelatin substrates, and the presence of transcellular pores is revealed using a fluorescent tracer (fluorescently labeled streptavidin). **b,** By fluorescence microscopy, transcellular pores are detectable as bright spots in the middle of dark islands that represent cells’ footprints. **c**, A step-by-step schematic of our mechanobiological assay tailored for transcellular pore induction and detection: **i)** Glass substrates were coated with biotinylated gelatin. **ii)** Particles were randomly seeded on the biotinylated gelatin for 10 minutes. **iii)** phSC cells were seeded over particles and cultured for 2-7 hours to fully spread. **iv)** The fluorescent tracer was added to media for 5 minutes to detect transcellular pores. Cells were then fixed and imaged. The particle is shown in grey or dotted circles (in top views of **iii** and **iv** when covered by the cell). **d**, Confocal images show significant cellular deformation localized at the particle site. Panels **i** and **ii** show top view, maximum intensity projections of a phSC cell labeled for cell cytoplasm and nucleus (red). The cell is spread above a particle (white arrow in panel **i**). The fluorescent label (shown in green; panel **ii**) is bound to the biotinylated gelatin substrate outside of the cell footprint. Panel **iii** shows a cross-sectional side view of the cell at the location of the particle (along the dashed line in panel **ii**) highlighting cell thinning above the particle (arrow). **e,** Estimated cell thickness from confocal images. Cells were significantly thinner above the particles (**** p < 0.0001). **f,** Estimated cell thickness from confocal images at nuclear and non-nuclear regions. **g**, Confocal images show both the cellular deformation and successful transcellular pore detection at the microparticle region (panels **i** and **ii**: top view, maximum intensity projections; panel **iii**: cross-sectional side view). A phSC cell is spread above a particle (white arrow in **i**). The fluorescent label is bound to the biotinylated substrate outside of the cell footprint and at the border of neighboring cells due to lack of tight intercellular junctions (**ii**). The fluorescent tracer is also bound to the substrate at the location of the particle indicating transcellular pore formation (highlighted by the arrow in **iii**). White arrow heads in panels **d** and **g** point to intracellular vesicles (see **Fig. S2**).

**Fig. 3c** illustrates the principle of the fluorescent assay to detect transcellular pores. High-definition confocal images of a representative cell overlying a Ø 5 µm particle are shown in **Fig. 3d**. After seeding over particles, cells naturally deformed, thinning directly above the particle (white arrow in panel iii inset). Cell thickness was quantified using cytoplasmic and nuclear staining in cross-sectional views, revealing that cell thickness above particles (1.6 ± 0.3 µm) was significantly less than beside the particles (5.9 ± 0.7 µm; two-tailed, paired t-test: p-value < 0.0001; **Fig. 3e**). We also measured cell thickness at nuclear (5.6 ± 1.1 µm) and non-nuclear (1.7 ± 0.4 µm) regions (**Fig. 3f**) which were comparable to cell thickness beside (p = 0.20) and above particles (p = 0.25), respectively.

**Fig. 3d** displays a thinned region of a phSC cell stretched over a particle. Notably, the fluorescent tracer did not adhere to the substrate near this particle (**Fig. 3d**, panels ii and iii), indicating the absence of pore formation. In contrast, in other cells (**Fig. 3g**), a cellular opening (transcellular pore) overlying the particle was observed, with the fluorescent tracer also adhering to the substrate, indicating transcellular pore formation. We also detected intracellular vesicles that appeared as empty spaces after cytoplasmic staining in our confocal images (white arrowheads in **Fig. 3d** and **Fig. 3g**). It is important to clarify that unlike transcellular pores, these vesicles are not openings extending through the cell and were also observed in cells that were not positioned over particles (**Fig. S2**).

### Transcellular pores are highly dynamic

To screen compounds that facilitate pore formation in cells, it is useful to increase the number of cells that are examined; further, it is essential to develop a protocol for testing the effects of investigational compounds on pore formation. Thus, we extended our original protocol to include a treatment step and a second tracer. In the modified protocol, cells were seeded over particles for 2-7 hours in 16-well chambered slides and exposed to a first fluorescent tracer for 5 minutes to detect particle-induced pores, as described above (**Fig. 4a**). Cells were then subjected to a “treatment” (here, an external magnetic field or a putative SC-active agent) for a defined time (10 minutes for the external magnetic fields and 30 minutes for the putative SC-active agent), after which they were incubated with a second fluorescent tracer for 5 minutes. We captured images using widefield microscopy (brightfield and fluorescence), which facilitated the examination of hundreds of cells per well (581 ± 189 cells/well).

**Fig. 4.**
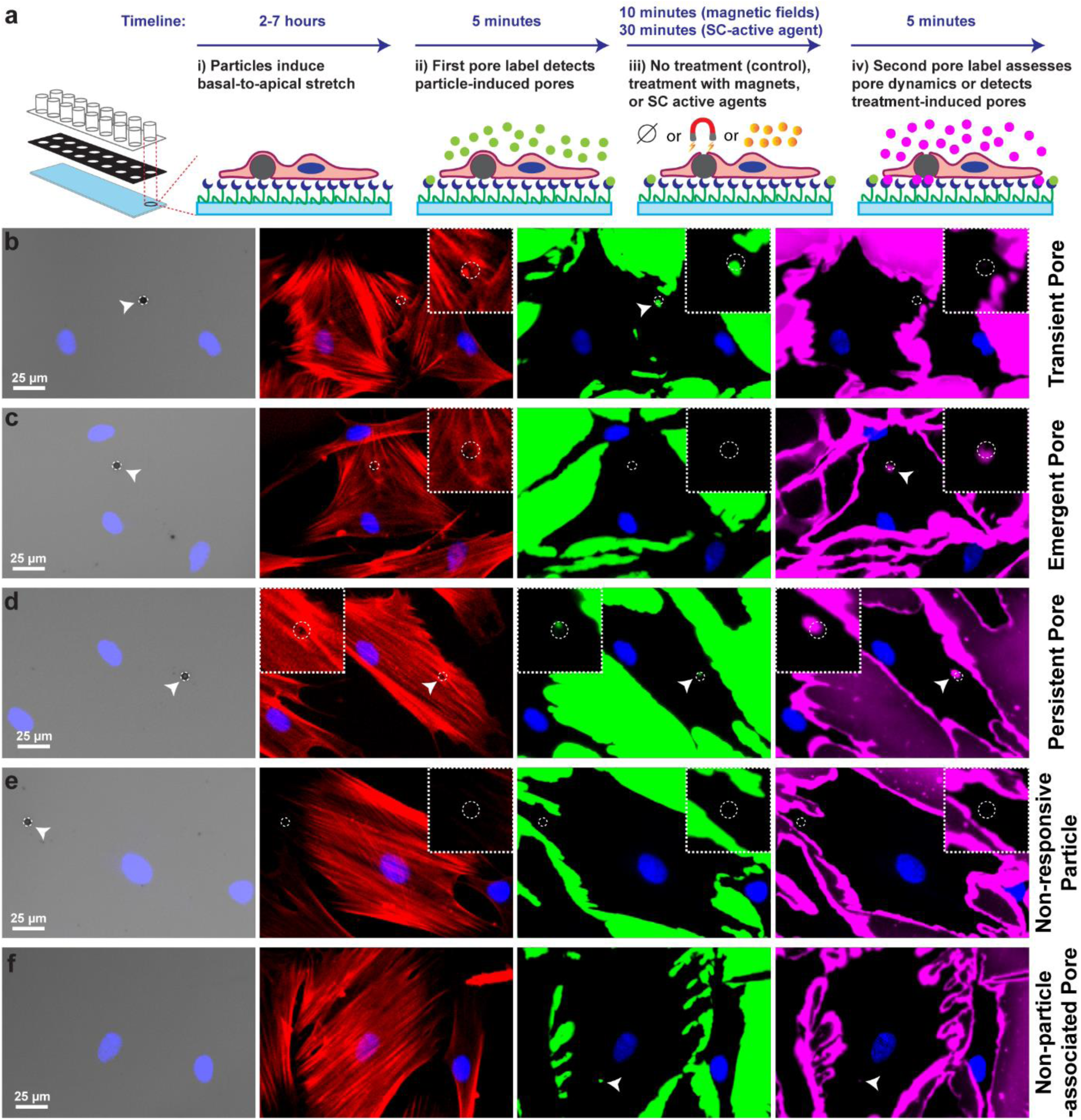
Fluorescence detection of different transcellular pore types. **a,** Schematics of the assay: phSC cells were cultured over particles on a biotinylated gelatin substrate in 16-well chambered slides to allow for high-content detection of transcellular pores. Initially, cells were exposed to a first fluorescent tracer (green) to detect any transcellular pores resulting from particle interaction alone. Subsequently, cells were treated with SC-active agents or exposed to external magnetic fields to induce further pore formation or left untreated as controls to assess pore dynamics. Pores induced by these treatments or new pores emerging in control conditions at particle locations were detected using a second tracer (magenta). **b,** Transient pore: A pore detected only by the first fluorescent tracer, indicating pore closure by the time of the second labeling step. **c,** Emergent pore: A pore detected only by the second fluorescent tracer, indicating it formed after the initial labeling step. **d,** Persistent pore: A pore detected by both the first and second fluorescent tracers. Note the rarefaction of the actin network at the pore location. **e,** Non-responsive particle: A particle without any detectable pores. **f,** Non-particle-associated pore: A pore detected by the fluorescent assay but not associated with any visible particle. Insets in panels **b-e** provide 2x magnified views at the location of particles. The first column in panels **b-f** show DAPI-stained nuclei (blue) merged with a brightfield image, where particles appear as black dots (white arrowheads) and are highlighted by dashed circles in other images. F-actin (red), first fluorescent tracer (green), and second fluorescent tracer (magenta) are shown in the second, third, and fourth columns of panels **b-f**, respectively.

We observed a diversity of pore formation behaviors, and for definiteness we classified pores based on their labelling patterns by the two fluorescent labels used in the assay:

- Transient pores were pores detected only by the first tracer, *i.e.*, they had closed at the time of the second labeling (**Fig. 4b**; see also “Persistent pores” below).
- Emergent pores appeared solely in response to the second tracer, indicating that they had formed after the first labeling period (**Fig. 4c**).
- Persistent pores were pores labelled by both tracers, *i.e.*, they were open at some point during both labeling steps (although not necessarily continuously; **Fig. 4d**). We note that persistent pores could appear to be transient if the biotin substrate were to be saturated by the first fluorescent tracer, thereby blocking labeling by the second tracer; however, even in regions of maximal labeling by the first tracer (see Methods), the second fluorescent tracer was typically detectable at the edges of the initial staining patterns, likely due to lateral diffusion. This enhances confidence in the true transient nature of pores identified as “transient pores”.

The existence of these different classes of pores indicates that the pore formation process is dynamic, and that pores can close in addition to opening in our assay. Here, the term ‘dynamic’ refers processes of pore formation and closure, which occurred within minutes to tens of minutes, *e.g.* transient pores were open during the initial 5-minute labeling period but were closed during the second 5-minute labeling period, which followed a 10-minute exposure to the magnetic field or a 30-minute exposure to a putative SC-active agent.

It is noteworthy that some particles did not induce any detectable pores with either label (noted as a non-responsive particle; **Fig. 4e**) and that we occasionally observed pores that were detected by the fluorescent tracer but were not associated with any particle (a non-particle-associated pore; **Fig. 4f**). These results underscore the complex nature of pore formation. It is also of interest to note that the spatial extent of second tracer labelling outside of and between cells was generally larger than the extent of labeling with the first tracer (**Fig. 4**), likely due to cell movement over the course of the assay and some lateral diffusion of the second tracer during its exposure time.

### Particles induce fewer transcellular pores in normal vs. glaucomatous phSC cells

SC pores are reduced in glaucoma,^23,24^ and we wondered whether this behavior would be replicated in our *in vitro* assay. We therefore compared particle-induced pore counts in phSC cell strains isolated from normal vs. glaucomatous donor eyes (**Fig. 5a**). Although we observed significant variability in particle-induced pore formation rate between the tested cell strains, pooling the data for normal and glaucomatous cell strains (**Fig. 5b**) indicated that the glaucomatous cell strains formed significantly fewer transcellular pores at particle locations vs. normal cells (7.0 ± 3.2 vs. 11.0 ± 5.0 % pore/particle; two-tailed Mann-Whitney test: p < 0.0001). Given that age is also a risk factor for glaucoma,^33–35^ we examined the influence of donor age on particle-induced pore formation rate in phSC cells. Interestingly, there was a correlation between particle-induced pore incidence and the age of normal phSC cell donors, but this correlation was weak (R^2^ = 0.10, p = 0.0006; **Fig. S3a**), suggesting that factors other than age also contribute significantly to the variability in pore numbers among normal phSC cells. When accounting for glaucoma status as an independent variable, multiple linear regression revealed a somewhat stronger correlation between particle-induced pore incidence rate and donor age across both normal and glaucomatous phSC cells (R^2^ = 0.39, p = 0.0004; **Fig. S3b**). Note that our glaucomatous cells were generally obtained from older donors compared to the normal cells (**Table 1**).

**Fig. 5.**
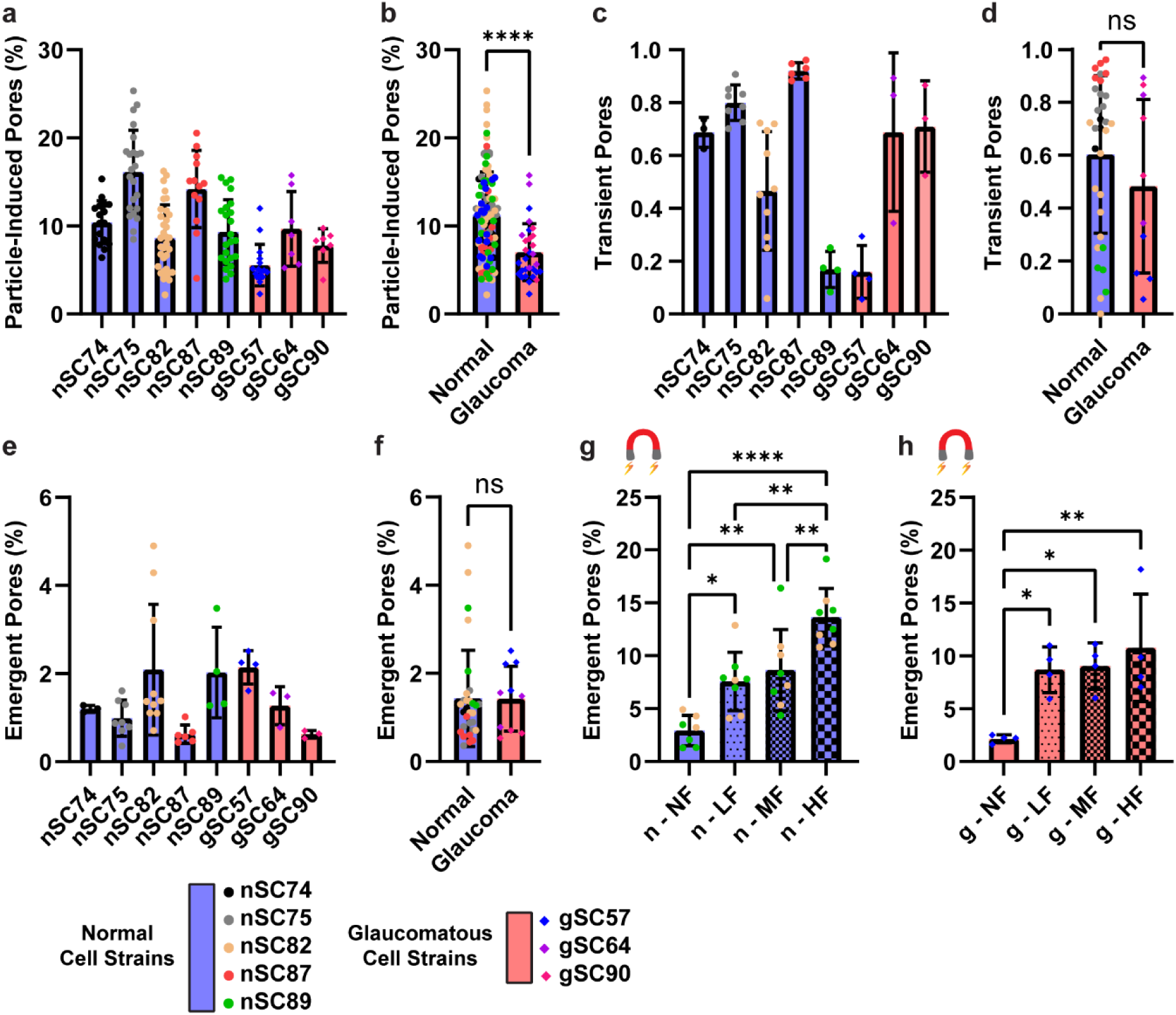
Analysis of particle-induced transcellular pores in normal and glaucomatous cell strains. **a,** Particle-induced transcellular pores detected by the first fluorescent tracer (“Particle-induced Pores”) in normal (blue bars, circular data points) and glaucomatous (orange bars, square data points) cell strains, presented as the number of pores divided by the total number of particles under cells, expressed as a percentage. The color and shape of data points for each cell strain remain consistent throughout the entire figure. **b,** Pooled data from panel (**a**) comparing particle-induced transcellular pores in normal and glaucomatous cell strains. **c,** Fraction of particle-induced pores in each cell strain that were detected by the first fluorescent tracer, as shown in panel (**a**), but were closed at the time of exposure to the second fluorescent tracer (“Transient Pores”). **d,** Pooled data from panel (**c**), comparing transient pores in normal and glaucomatous cell strains. Note the values in panels **c** and **d** are the fraction of particle-induced pores that were transient, which differs from the interpretation of percentages in other panels. **e,** Percentage of particles that induced new pores detected only by the second fluorescent tracer (“Emergent Pores”). **f,** Pooled data from panel (**e**), comparing emergent pores in normal and glaucomatous cell strains. **g** and **h,** Percentage of emergent pores detected only by the second fluorescent tracer in normal (panel **g**) and glaucomatous (panel **h**) cell strains exposed to no force (NF; control), low force (LF), medium force (MF), or high force (HF) conditions. n: normal; g: glaucomatous. See **Fig. S4** for magnetic force application and measurement. Data are presented as mean ± standard deviation. Statistical significance is indicated with (ns), (*), (**), (***), and (****) corresponding to not significant, p < 0.05, p < 0.01, p < 0.001, and p < 0.0001, respectively, except in panels **a**, **c**, and **e**. Total numbers of pores counted in each of normal and glaucomatous cells for panels **b**, **d**, **f**, **g**, and **h** are indicated in **Table S1**.

**Table 1.**
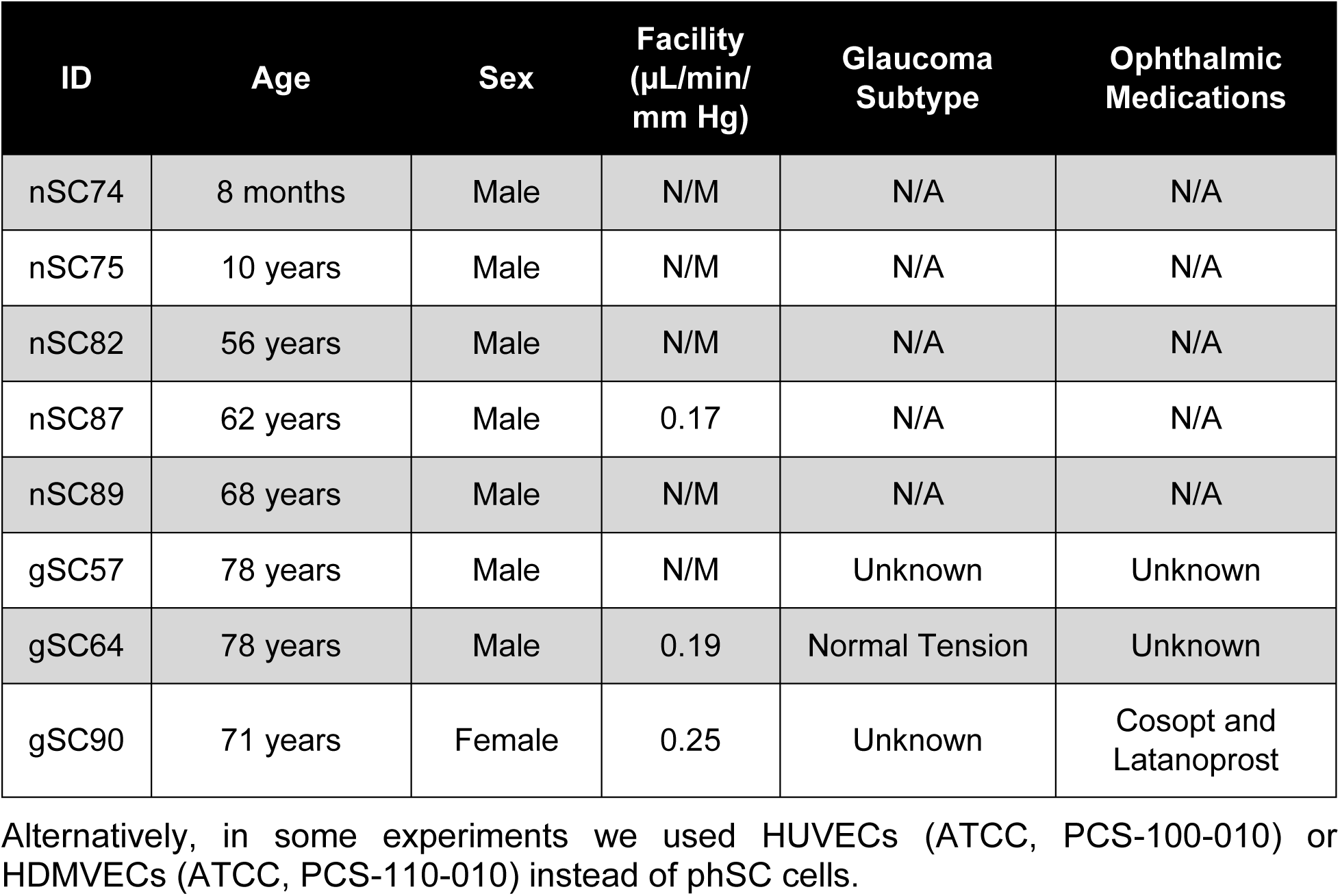
Summary of cell strains. The initial letter in the cell strain ID indicates whether the donor had a history of glaucoma (g) or if the cells were obtained from normal (n) eyes. N/M = not measured. N/A = not applicable.

| ID | Age | Sex | Facility<br>($\mu\text{L}/\text{min}/\text{mm Hg}$) | Glaucoma<br>Subtype | Ophthalmic<br>Medications |
| --- | --- | --- | --- | --- | --- |
| nSC74 | 8 months | Male | N/M | N/A | N/A |
| nSC75 | 10 years | Male | N/M | N/A | N/A |
| nSC82 | 56 years | Male | N/M | N/A | N/A |
| nSC87 | 62 years | Male | 0.17 | N/A | N/A |
| nSC89 | 68 years | Male | N/M | N/A | N/A |
| gSC57 | 78 years | Male | N/M | Unknown | Unknown |
| gSC64 | 78 years | Male | 0.19 | Normal Tension | Unknown |
| gSC90 | 71 years | Female | 0.25 | Unknown | Cosopt and<br>Latanoprost |

We further investigated the pore dynamics in normal and glaucomatous cells by counting the number of pores detected by the first fluorescent tracer which were closed 10 minutes later when the cells were exposed to the second fluorescent tracer without any treatment (transient pores; **Fig. 5c**). Pooling the data for normal and glaucomatous cells (**Fig. 5d**) indicated that the fraction of particle-induced pores that were transient was not significantly different between normal and glaucomatous cells (0.60 ± 0.30 vs. 0.48 ± 0.33 transient pore/particle-induced pore; two-tailed Mann-Whitney test: p-value = 0.3706), suggesting that the duration of pores is not markedly different between normal and glaucomatous phSC cells.

We also examined the incidence of emergent pores, *i.e.*, pores detected by the second fluorescent tracer but not by the first fluorescent tracer without any treatment (**Fig. 5e**). Pooling the data (**Fig. 5f**) showed that the percentage of emergent pores was also not significantly different between normal and glaucomatous cells (1.4 ± 1.1 vs. 1.4 ± 0.7 % pore/particle; two-tailed Mann-Whitney test: p-value = 0.5053).

### Magnetic force enhances transcellular pore formation in normal and glaucomatous phSC cells

We applied an external magnetic field to the magnetic microparticles in the dish, with the expectation of promoting cell stretching, thus reducing the thickness of overlying cells and increasing pore formation rate above that seen when cells are simply seeded on particles without magnetic force application. Following the application of the magnetic field (see **Fig. S4**), we observed more pores in both normal and glaucomatous cells (**Fig. 5g** and **Fig. 5h**), even at the weakest magnetic field strength tested (a force of 235 ± 9 pN per particle). Specifically, at this low force level, there were 2.6-fold more magnet-induced pores in normal cells (from 2.9 ± 1.4 to 7.6 ± 2.8; ANOVA p = 0.0200) and 4.1-fold more for glaucomatous cells (from 2.1 ± 0.4 to 8.7 ± 2.2; ANOVA p = 0.0392) vs. the control group without magnetic force application. Similarly, at the highest magnetic field strength tested (461 ± 10 pN per particle), there were 4.7-fold more magnet-induced emergent pores in normal cells (from 2.9 ± 1.4 to 13.6 ± 2.7; ANOVA p < 0.0001) and 5.0-fold more in glaucomatous cells (from 2.1 ± 0.4 to 10.8 ± 5.1; ANOVA p = 0.0070) vs. the control group. We conclude that magnetic force delivery enhances pore formation in a field strength-dependent manner in our assay.

### Pore formation is strongly associated with native and modified phSC cell stiffness

It is known that the inner wall of SC is stiffer in glaucomatous eyes with elevated flow resistance vs. in normal eyes.^36^ Based on this observation, we wondered whether pore formation was correlated with cell stiffness. We therefore measured stiffness in untreated cells using atomic force microscopy (AFM) on five normal and 3 glaucomatous cell strains (**Fig. 6a**). A regression analysis revealed a significant correlation between particle-induced pore incidence and cell stiffness (R^2^ = 0.41, p < 0.0001; **Fig. 6b**).

**Fig. 6.**
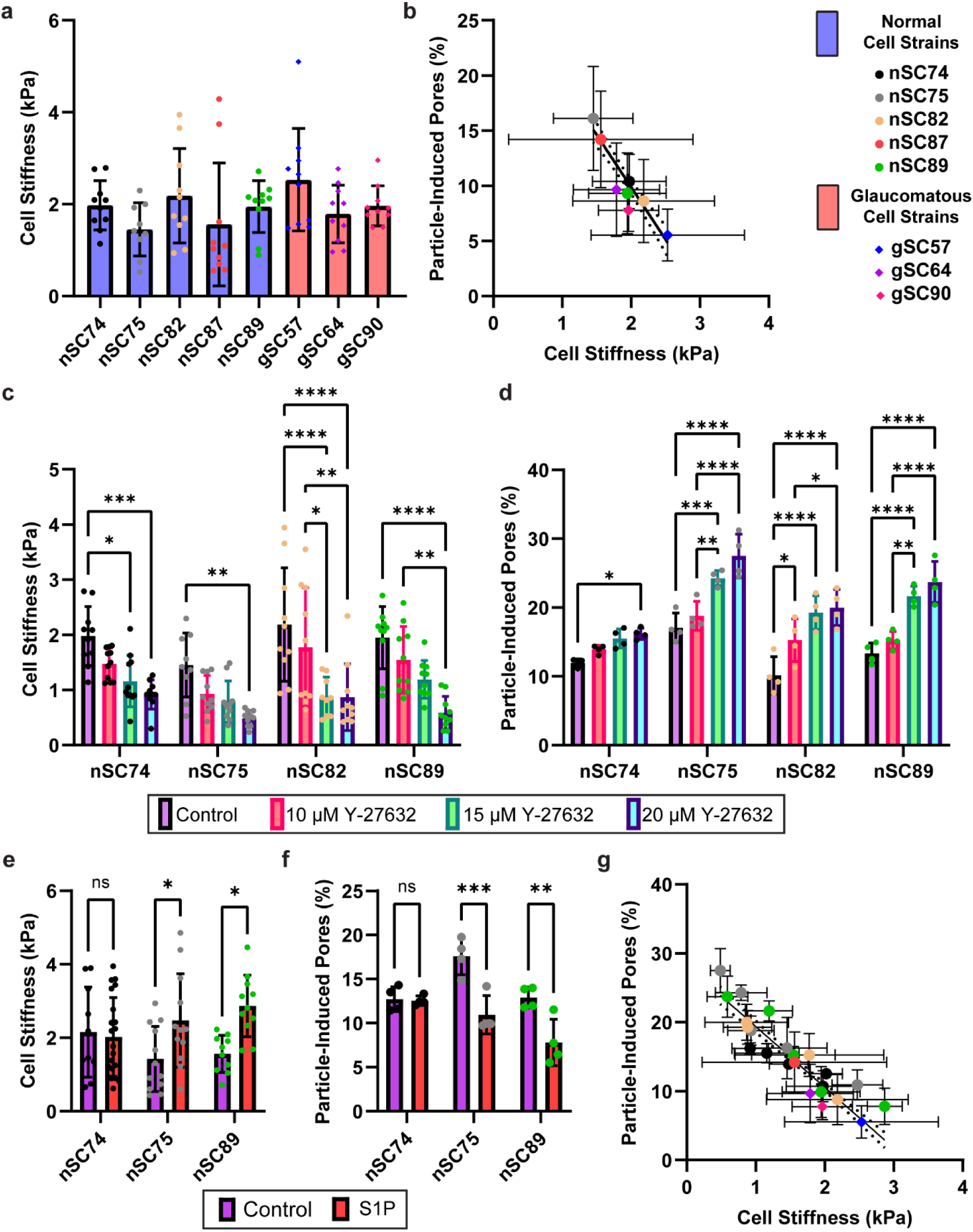
The impact of cell stiffness on pore formation. **a,** Stiffness of five normal cell strains and three glaucomatous cell strains measured by AFM. **b,** Linear regression of cell stiffness and particle-induced pore incidence (R^2^ = 0.41). Dotted lines show 95% confidence intervals. **c,** Cell stiffness was decreased by Y-27632 treatment in a dose-dependent manner in four normal cell strains, while **d** pore formation was increased by Y-27632 treatment in a dose-dependent manner in these same strains. Color legend is the same in panels **c** and **d**. **e** and **f,** Impact of S1P on cell stiffness and pore formation. S1P treatment (0.1 μM; 30 minutes) resulted in a significant increase in cell stiffness **(e)** and a decrease in particle-induced pore formation **(f)** in two phSC cell strains, nSC75 and nSC89. However, S1P treatment did not affect cell stiffness or pore formation in strain nSC74. Color legend is the same in panels **e** and **f**. **g,** Linear regression of particle-induced pore incidence on cell stiffness after Y-27632 or S1P treatments in four normal cell strains (nSC74, nSC75, nSC82, nSC89) and in all untreated SC cell strains (R^2^ = 0.63) The color legend is the same in panel **a** and in **b**, noting that points with the same color in panel **g** refer to a single cell strain under both untreated and treated conditions. Data are presented as mean ± standard deviation. Statistical significance is indicated with (ns), (*), (**), (***), and (****) corresponding to not significant, p < 0.05, p < 0.01, p < 0.001, and p < 0.0001, respectively.

We then asked whether, within a given cell strain, modulation of cell stiffness would affect pore formation. We first lowered cell stiffness using a Rho-associated kinase inhibitor (ROCKi) Y-27632. Such compounds are particularly relevant to glaucoma, since two ROCKi compounds have been approved for clinical reduction of IOP, primarily by reducing AH outflow resistance. Further, ROCK inhibition has been shown to decrease actin stress fibers in SC cells.^37,38^ We found that Y-27632 decreased cell stiffness (**Fig. 6c**) and increased pore formation in a dose-dependent manner in all tested phSC cell strains (**Fig. 6d**). Note that the Y-27632 treatments did not significantly affect the cell area (**Fig. S5**).

We next asked whether the converse effect would occur, *i.e.*, would cell stiffening decrease pore formation rate within a given cell strain. For this purpose we used sphingosine-1-phosphate (S1P), known to increase cell contractility^39^ and stiffness^21,40^ of phSC cells, and to elevate flow resistance of the conventional outflow pathway.^41^ We found that S1P increased cell stiffness and decreased pore formation in the nSC75 and nSC89 cell strains yet did not have a significant effect on cells from the nSC74 strain (**Fig. 6e** and **Fig. 6f**). Importantly, regression analysis unveiled a robust correlation between particle-induced pore incidence rate and cell stiffness (R^2^ = 0.63, p < 0.0001). This correlation included natural stiffness variations between untreated SC cells strains, as well as the stiffness-altering effects of Y-27632 and S1P across four cell strains.

### Particle-induced transcellular pore formation is directional

phSC cells experience a biologically rare basal-to-apical deformation *in vivo* and we show here that seeding cells above particles, mimicking certain aspects of the *in vivo* deformation experienced by SC cells, induces transcellular pores *in vitro* (**Fig. 2**, **Fig. 3**, and **Fig. 4**). Interestingly, when we placed particles <u>above</u> phSC cells to apply an apical-to-basal local cellular deformation, we failed to detect any transcellular pores using our fluorescent assay at the location of particles (**Fig. S6** and **Fig. S7**). Even applying an apical-to-basal magnetic field (473 ± 11 pN per particle; see Methods and **Fig. S4** for magnetic force measurement) or treatment with 10 µM Y-27632 did not induce transcellular pores. This shows that transcellular pore formation in SC cells is “direction dependent.”

In diapedesis, leukocytes move directly through individual endothelial cells by forming apical to basal transcellular routes,^5,42^ which is opposite to the direction of AH transport through pores formed in SC endothelial cells. Consequently, we posited that positioning particles atop cells capable of facilitating leukocyte transfer from the apical to basal direction *in vivo* would prompt the formation of transcellular pores in our *in vitro* assay. We therefore seeded particles atop human umbilical vein endothelial cells (HUVECs) and human dermal microvascular endothelial cells (HDMVECs). Intriguingly, we routinely observed the formation of transcellular pores at the sites where particles were seeded on top of both HUVECs and HDMVECs (see **Fig. S8**), which was dramatically different to that seen in SC cells.

## Discussion

We here describe the design and implementation of an *in vitro* platform that applies relevant basal-to-apical local cell stretching to create transcellular pores in adherent cells, with particular focus on transcellular pores in phSC cells. By quantifying pores in both normal and glaucomatous phSC cells, we demonstrate that our platform replicates important aspects of *in vivo* pore formation, as discussed below, and provides a useful tool for studying basic mechanisms of transcellular pore formation. With further development, we expect that this platform could be used for high content screening of compounds that can modulate pore formation, of great interest in the management and treatment of ocular hypertension.

Our *in vitro* assay successfully replicated important *in vivo* mechanobiological observations. For example, glaucomatous SC cells exhibited impaired particle-induced pore formation, consistent with previous observations of reduced pore forming ability in glaucomatous cells in native tissue (and in previous *in vitro* assays based on substrate stretching or transcellular perfusion).^20,21^ Our assay also demonstrated the dynamic nature of pores, expected because of the dynamic mechanobiological environment within the outflow pathway,^43^ and the tendency of pores to form at locations of cell thinning over particles, consistent with the tendency of pores to form in association with the thinnest regions of giant vacuoles *in vivo*.^44^ Further, we observed a correlation between the focal force delivered magnetically and the formation of pores, consistent with pore formation being a mechanobiologically-mediated process.^21^

The key finding of this work is that there is a strong association between cell stiffness and transcellular pore forming ability. This association was present when comparing cell strains with naturally occurring stiffness differences and when modulating cell stiffness with cell softening or stiffening agents, namely the ROCK inhibitor Y-27632 and S1P, respectively. Importantly, this association is consistent with the diminished pore formation ability of glaucomatous cells and reinforces the idea that SC cell stiffness is a fruitful therapeutic target for IOP lowering in glaucoma patients. For example, targeted latrunculin A micelles have been shown to reduce IOP via softening SC cells in an animal model.^45^ Currently, no approved medications have been shown to directly target SC pore formation, but some, such as the ROCKi Netarsudil,^46^ may affect SC pores as part of their mode of action. Interestingly, the pore-forming response to magnetic force was higher in glaucomatous cells compared to normal cells. When a minimal force of ~250 pN per particle was applied, corresponding approximately to a local pressure of ~0.1 mmHg over the surface of the particle, there was a significant 4.1-fold increase in pore formation for glaucomatous cells compared to only 2.6-fold increase for normal cells. Although many possible explanations exist for this finding, these results hold out hope that impaired pore formation in glaucomatous cells could be rescued by further stretching and thinning of these cells. This raises important questions: Is there a minimum cell thickness that needs to be achieved to initiate pore formation? Could normal SC cells be more agile or dynamic in accommodating forces compared to glaucomatous cells? Is there an upper limit to the number of pores that a cell can form at the location of each GV/particle?

The obvious question is a mechanistic one: why do softer cells more readily form pores? We know that pore formation is triggered by mechanical stimulation (stretch),^20^ and our SEM micrographs show pores forming at the location of particles where the cell membrane is thinned. Because pore formation requires near-apposition and fusion/reorganization of the basal and apical cell membranes, it seems likely that the key step in pore formation is cell thinning, and that this thinning process is facilitated in softer cells. We thus propose an *in vivo* model of transcellular pore formation in phSC cells, where a relatively soft normal phSC cell experiences significant stretching at the location of GVs and consequent transcellular pore forms at the thinnest location of the cell associated with the GV (**Fig. 7a**). In contrast, the stiffer glaucomatous cells do not thin enough to initiate pore formation (**Fig. 7b**). Our model thus suggests the testable hypothesis that there is a minimum cell thickness that needs to be achieved to initiate pore formation.

**Fig. 7.**
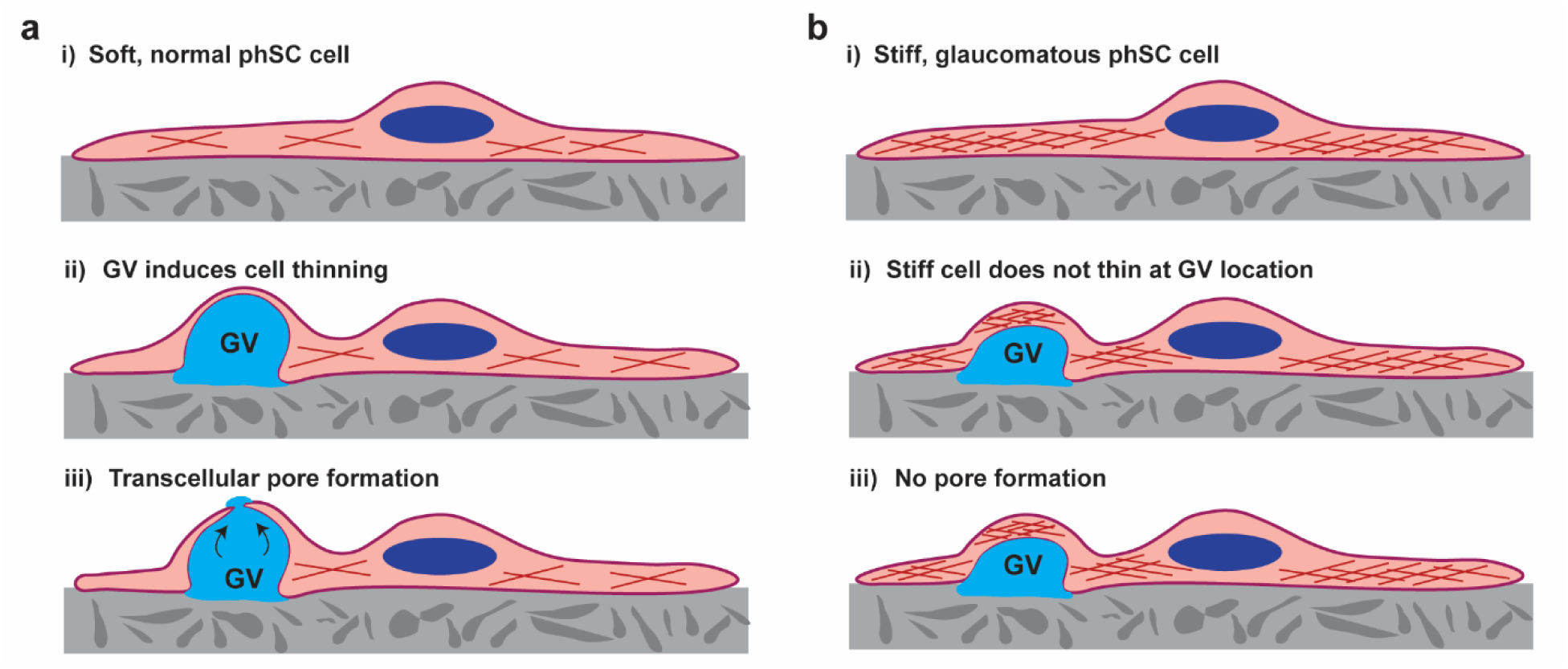
Proposed model of intracellular pore formation in phSC cells. **a,** In a normal phSC cell (**i**), GV formation results in substantial cell thinning (**ii**), eventually causing cell membranes to come into close proximity and induce pore formation. **b,** In a stiff glaucomatous cell (**i**), GV formation is unable to sufficiently stretch and thin the cell (**ii**), preventing the cell membranes from coming close enough to facilitate pore formation (**iii**). Cytoskeleton filaments, such as actin filaments, are depicted in red, cell nuclei in dark blue, AH in light blue, and both the extracellular matrix and BM in grey.

Here, it is important to note that a putative mechanistic link between a softer cell and more cell stretching/thinning, while appearing simple, actually hides significant complexity. Notably, many cellular biophysical properties are associated with cell stiffness, including the fact that stiffer cells exert more traction force on their substrate.^47,48^ This would suggest that stiffer cells overlying beads would generate <u>more</u> traction-induced deformation and cellular thinning over the bead, which according to our understanding, would increase, rather than decrease, pore formation. Clearly the situation would benefit from future studies of additional biomechanical readouts, such as those offered by traction force microscopy.^49,50^ Further, our AFM measurements were made using a 10-micron tip over a time scale of seconds, so that the measured properties represent the acute mechanical response of the cell’s sub-cortical cytoskeleton.^21^ Conversely, our assay allows cells to spread over microparticle for hours, during which time the cell presumably remodels its cytoskeleton and adapts its physical configuration to accommodate the presence of the microparticle. It thus could be argued that it is somewhat surprising that a correlation between AFM-measured cell stiffness and pore-forming ability exists. Nonetheless, there is a strong association, and this intriguing finding strongly motivates future mechanistic studies.

In *ex vivo* settings, it has been shown that SC pores are reduced in glaucoma.^23,24^ Here, as well as in previous work,^21^ we demonstrated that this same behavior is preserved in SC cells *in vitro*, further validating their use as a model for glaucoma. An additional observation that supports the use of these cells as a robust *in vitro* model is the directional dependence of pore formation. In our assay, we observed that basal-to-apical force induces pore formation in SC cells, whereas apical-to-basal force does not. This unique mechanobiological behavior of SC cells *in vitro* mirrors their specialized function *in vivo*, providing further motivation for using these cells to model the mechanophysiological properties of the outflow pathway in glaucoma.

Naturally, our work is subject to several constraints. The first relates to the time required for cell spreading over microparticles. The phSC cells exhibited a spreading duration ranging from 2 to 7 hours, with variations observed among different cell strains. Prolonged culturing times increase the risk of microparticle endocytosis, remodeling of the gelatin coating, and potential disruption of the assay. Consequently, optimizing the duration that phSC cells remain in contact with microparticles and the gelatin substrate is crucial. Alternatively, microparticle surface modifications may minimize the risk of microparticle endocytosis, although in our hands PEGylation, usually protective against internalization, led to endocytosis in our assay. For this reason, all assays were carried out with the beads as supplied.

Lastly, the quantification of pores was constrained by the need for manual counting (*i.e.*, identifying fluorescent signals at the particle locations). Pore identification is challenging for multiple reasons, especially when cells were densely seeded. First, since phSC cells do not form tight junctions *in vitro*, fluorescent tracer can adhere to the substrate in a “patchy” pattern at cell-cell margins, generating signals that are hard to distinguish from pores. For this reason, we focused exclusively on transcellular pores, excluding paracellular pores. Additionally, the lateral diffusion of the fluorescent tracer under cells (at pore locations) and/or cell movement can cause pores to appear larger than their actual size. The accuracy of the fluorescent assay could be enhanced by developing methods that enable the formation of tight junctions by phSC cells. Additionally, implementing computer-aided algorithms for pore identification would streamline and improve the overall process.

In summary, this study presents a novel and valuable tool for understanding transcellular pore formation. We find that pore forming ability is strongly associated with cell stiffness in phSC cells. We expect that by elucidating the mechanisms underlying pore formation in SC cells, we can accelerate the development of innovative treatments targeting ocular hypertension in glaucoma patients.

## Methods

### Isolation and characterization of phSC cells

To account to human variability, a total of five phSC cell strains isolated from five healthy donor eyes and three cell strains isolated from three donors having a history of glaucoma were used in this study (**Table 1**). Use of human donor tissue adhered to the tenets of the Declaration of Helsinki. phSC cells were harvested using a cannulation method in human donor eyes, and were then cultured and characterized as previously described.^25,26^ Characterization was based on the cell’s typical spindle-like elongated cell morphology, expression of vascular endothelial-cadherin and fibulin-2, a net transendothelial electrical resistance of 10 ohms·cm^2^ or greater, and the absence of myocilin induction following exposure to dexamethasone. The SC cells were cultured in low glucose Dulbecco’s Modified Eagle Medium (DMEM; Gibco 11885084) supplemented with 10% fetal bovine serum (FBS), and 1x penicillin streptomycin glutamine (PSG; Gibco 10378016) at 37°C with 5% CO_2_.

### Gelatin biotinylation

A gelatin solution (Type B, 1% in H_2_O, Sigma-Aldrich G1393) was dialyzed against 0.1 M NaHCO_3_ at pH 8.3. The biotinylation reagent (EZ-Link™ NHS-LC-LC-Biotin, Thermo Scientific 21343, in dimethyl sulfoxide) was then mixed in 12-fold excess molarity with the gelatin solution at room temperature for 1 hour. Subsequently, the biotinylated gelatin solution underwent dialysis against phosphate buffer saline (PBS) to remove excess biotin. The purified biotinylated gelatin was aliquoted and stored at 4°C or preserved at −20°C for long-term storage.

### Substrate preparation and cell culture for transcellular pore formation using microparticles

Round cover glasses (12 mm in diameter, #1.5 Thickness; Harvard Apparatus, 64-0718) were employed for SEM imaging, while 16-well chambered cover glasses (16 wells per cover glass; Invitrogen C37000; **Fig. 4 A**) were utilized for other experiments. Glass substrates underwent plasma treatment (Harrick Plasma, PDC-32G) for 2 minutes at high RF power and were immediately incubated at 37°C with 0.5 mg/mL biotinylated gelatin for 24 hours. After three washes with PBS, the glass substrates were incubated with microbial transglutaminase (0.001 unit/mL in 20 mM Tris-HCl and 150 mM NaCl, pH 8; Sigma-Aldrich, SAE0159) at 37°C for 24 hours to crosslink the gelatin coating and enhance phSC cell attachment on the substrate. Following three additional washes with PBS, the cover glasses were exposed to UV light in a biosafety cabinet for 1 hour to sterilize the substrate and denature the transglutaminase. Carboxyl ferromagnetic particles (4.0-4.9 µm; Spherotech Inc., CFM-40-10) were vortexed and added to the substrate. The particle size was based on the volume of GVs with transcellular pores. Assuming a spherical shape, the diameter of GVs, as reported by Swain *et al*.,^44^ ranged from 4.8 to 7.5 µm. The particles were allowed to randomly sink onto the substrate to achieve an approximate density of 1 particle per 1000 µm^2^. Non-adherent particles were removed by washing three times with PBS, followed by seeding phSC cells (passage 4-6; 7.5 k cells/cm^2^). The cells were then cultured for 2 to 7 hours in DMEM supplemented with 1% FBS and 1x PSG at 37°C with 5% CO_2_. The culture time was chosen as the minimum duration necessary for the cells to adopt a spread configuration similar to what observed *in situ*.^22^ Note that this culture time (2 to 7 hours) varied between the tested cell strains. Alternatively, in some experiments, apical-to-basal cell stretching was achieved by adding particles above fully spread cells.

### SEM sample preparation and imaging

Cells were fixed in universal fixative (2.5% glutaraldehyde, 2.4% paraformaldehyde, and 0.08 M Sorensen’s phosphate buffer in PBS) for 30 minutes and washed three times with PBS. Following fixation, the cells were incubated in a 2% tannic acid and guanidine hydrochloride solution in the dark for 2 hours, and then washed three times in PBS over a 60-minute period. Subsequently, the cells were post-fixed in 1% osmium tetroxide in PBS for 60 minutes, washed three times with PBS, and then rinsed three times in ultra-pure water. The cells were then dehydrated in an ethanol series (25, 50, 75, 90, 100, 100, and 100%; 10-minute changes), transferred to a 1:1 mix of 100% ethanol:hexamethyldisilazane (HMDS) for 10 minutes, followed by two 10-minute changes in HMDS, and air-dried in a fume hood with vigorous shaking. Further drying of the cells took place overnight in a chemical hood. Subsequently, the cells were mounted on SEM stubs with conductive carbon adhesive tape. Samples were coated with approximately 10 nm of gold/ platinum using a spotter coater and examined with a Hitachi SU8010 Cold Field Emission SEM.

### Pore detection using fluorescent assay and confocal microscopy

Once cells had fully spread over particles, they were incubated with the first fluorescent tracer (5 μg/mL Streptavidin, Alexa Fluor 488 Conjugate; Invitrogen, S32354) for 5 minutes at 37°C. Subsequently, the cells were immediately washed three times with PBS, fixed with 4% formaldehyde for 15 minutes at room temperature, washed three times with PBS, permeabilized with 0.1% Triton X for 15 minutes, and stained with 10 μg/mL HCS CellMask Stains (Invitrogen, H32712) for 1 hour. Following these steps, cells were washed three times with PBS and stored in PBS at 4°C until examination with a Zeiss LSM 900 confocal microscope system.

### Transcellular pore induction via magnetic force delivery

Upon achieving full cell spreading over particles, cells were incubated with the first fluorescent tracer for 5 minutes, followed by three washes with cell media. Subsequently, cells were either exposed to a magnetic field (as detailed below) or incubated solely with cell media for 10 minutes. Afterward, the cells were incubated with the second fluorescent tracer (5 μg/mL Streptavidin, Alexa Fluor 647 Conjugate; Invitrogen, S32357) for 5 minutes, all at 37°C. The cells were then immediately washed three times with PBS, fixed with 4% formaldehyde for 15 minutes at room temperature, washed three times with PBS, permeabilized with 0.1% Triton X for 15 minutes, and stained with NucBlue Fixed Cell Ready Probes Reagent (6 drops/mL; Invitrogen, R37606) and Alexa Fluor 555 Phalloidin (0.165 μM; Invitrogen, R34055) for 1 hour at room temperature. Following these steps, cells were washed three times with PBS, the cover glass was detached from the rest of the 16-well chambered system, mounted on another cover glass using Prolong Gold Antifade Mountant (Invitrogen, P10144), and stored in the dark until examination with a Leica DM6 B upright microscope system.

### Magnetic field delivery and measurement

Magnetic microparticles were exposed to a magnetic field for 10 minutes between the first and second pore labeling steps. The 10-minute exposure time was derived from live-cell imaging studies of lymphocyte diapedesis on endothelial monolayers by Carman *et al*.,^5^ who reported that the total duration of diapedesis ranged from approximately 4 to 8 minutes. Additionally, lymphocytes spent about 1 to 2 minutes at a location before initiating diapedesis. Utilizing a custom-designed magnetic actuator, neodymium N52 cylindrical magnets (3/16” diameter x 1/2” thick; D38-N52, K&J Magnetics) were precisely positioned in each well of the 16-well chambered slides at approximately 0.20 mm (high force; estimated 461 ± 10 pN on each particle), 0.55 mm (medium force; 337 ± 11 pN), or 0.91 mm (low force; 235 ± 9 pN) above the cells (**Fig. S4**). These magnetic forces are physiologically relevant, as the net force acting in the basal-apical direction of a GV is approximately 600 pN.^22^ Alternatively, in experiments where magnetic particles were situated atop cells to induce apical-to-basal local cell deformation, D38-N52 magnets were arranged within a well-plate, with the 16-well chambered slides directly positioned atop the magnets. In this setup, magnets were separated from the particles by a distance of 0.17 mm (equivalent to the thickness of the #1.5 coverslip), generating a magnetic force of 473 ± 11 pN per particle.

As shown in **Fig. S4**, the magnetic force exerted on individual particles was assessed by positioning magnets within a well-plate atop a calibrated scale, with the scale zeroed. Employing a high-precision Z-axis stage (Standa; Motorized Vertical Translation Stage, 8MVT188-20), a coverslip was delicately situated slightly above the magnets, ensuring no direct contact. A known quantity of particles, based on the particle concentration provided by the supplier and pipetting a known volume of particle solution, was then pipetted above the magnets, and the scale was read to determine the aggregate axial force applied on the particles. Through incremental adjustments to the distance between the magnets and particles utilizing the Z-axis stage, the magnetic force acting on particles at various distances from the magnet surface was measured (n = 3 repeats). Given that the magnetic force opposed the gravitational force on the magnets registered by the scale, alterations in the scale reading were utilized to determine the total magnetic force applied to the particles/magnets at specified distances. The magnetic force exerted on each particle was subsequently calculated by dividing the total force by the number of particles.

### Treatment with Y-27632 and S1P

Cells were seeded above microparticles in 16-well chambered biotinylated-gelatin coated slides in DMEM supplemented with 10% FBS, 1x PSG and allowed to spread for 3.5 hours at 37°C with 5% CO_2_. Subsequently, the cells were incubated with 10 μM, 15 μM, or 20 μM Y-27632 (Sigma, Y0503) or 0.1 μM S1P (TOCRIS, #1370) for 30 minutes, followed by the fluorescent tracer reagent with the treatments for 5 minutes. A 30-minute Y27632 treatment has been shown to be effective in reducing transforming growth factor beta2 (TGFβ2)-induced F-actin and α-smooth muscle actin (αSMA) upregulation.^37^ Following incubation, the cells were washed, fixed, stained for nuclei and actin filaments, and imaged as described below. We also determined the effects of Y-27632 on cell area. In brief, using FIJI software (NIH, Bethesda, MD, USA), cellular domains were determined by generating binary masks using thresholding of fluorescent actin images. Cellular masks were then used to calculate cell spread area. The average area per cell was then determined by dividing the total cell area within a defined region of interest (2 mm x 2 mm) by the total number of cells, based on number on nuclei in the same area. At least 400 cells were analyzed from each of the 4 normal phSC cell strains with 4 replicates per cell strain.

### Fluorescent imaging and analysis of transcellular pore incidence

For samples prepared on 16-well chambered cover glasses, fluorescent and brightfield images were acquired using a 10x objective to generate tile scans for each well. To avoid edge artifacts due to uneven gelatin coating, we restricted analysis to the cells within a 4 mm diameter circle centered in each well. The number of cells, particles beneath cells, and pores were manually counted. The percentage of pore incidence was quantified as the number of pores associated with particles divided by the total number of particles, multiplied by 100.

To determine whether a particle induced a pore, we required there to be a discrete fluorescently labeled region at a particle location. To determine the presence of labeling, we set a lower detection threshold for each fluorescent channel to be twice the average background intensity, as measured as the mean fluorescence signal from fully cell-covered regions. A signal above this threshold at the particle’s location was considered indicative of a pore.

Occasionally, we observed that the first fluorescent tracer appeared to saturate, or maximally label, the substrate, which could interfere with the detection of subsequent labeling by the second tracer. We defined “maximal labeling” by the first tracer as the condition where the intensity of the second label in areas without cells matched the background intensity in fully cell-covered regions. The presence of this labeling pattern would suggest that any additional binding sites available for labeling had already been occupied by the first tracer. However, even under maximal labeling conditions, the second tracer typically produced detectable signals at the edges of initial staining patterns, likely due to lateral diffusion of the fluorescent molecules, enabling us to conclude that the pore was open during exposure to the second tracer.

### Cell stiffness measurement

phSC cells (passage 4-6) were seeded on gelatin (0.5 mg/ml)-coated coverslips (12 mm in diameter; Cellvis, #SL39301002) at a density of 7.5 k cells/cm^2^ in low glucose DMEM supplemented with 10% FBS and 1x PSG and maintained at 37°C with 5% CO_2_ overnight. Subsequently, the cells were incubated with 10 μM, 15 μM, or 20 μM Y-27632 in or 0.1 μM S1P for 30 minutes, followed by cell stiffness measurements. An MFD-3D AFM (Asylum Research, Santa Barbara, CA, USA) was used to make stiffness measurements using silicon nitride cantilevers with an attached borosilicate sphere (diameter = 10 μm; nominal spring constant = 0.1 N/m; Novascan Technologies, Inc., Ames, IA, USA). Cantilevers were calibrated by measuring the thermally induced motion of the unloaded cantilever before measurements. The indentation depth was limited to 400 nm to avoid substrate effects and the tip velocity was adjusted to 800 nm/s to avoid viscous effects.^51^ 5 measurements/cell were conducted, and at least 10 cells were measured per condition. Data from AFM measurements were fitted to the Hertz model with setting the Poisson’s ratio is 0.5 to calculate the effective Young’s Modulus of the cells.

### Statistical analysis

Statistical analysis was conducted using GraphPad Prism, with significance set at p < 0.05. Descriptive statistics were calculated as mean and standard deviation. Data sets were evaluated for normality using the Shapiro-Wilk normality test. Those data sets passing the normality test, we carried out statistical comparisons using t-tests or ANOVA followed by Tukey’s multiple comparison test, as described. In cases where normality was not met, the non-parametric Mann-Whitney test or Kruskal-Wallis test followed by Dunn’s multiple comparisons test were performed. Significance levels in figures are denoted by (ns), (*), (**), (***), and (****) for not significant, p < 0.05, p < 0.01, p < 0.001, and p < 0.0001, respectively.

## Acknowledgements

BrightFocus Foundation: CG2020001 (CRE), G2023011S (DRO), G2021005F (BNS); National Institutes of Health: 5R21EY033142-02 (CRE and DRO), R01EY028608 (WDS), R01EY022359 (WDS and DRO), K99EY035360 (BNS), T32 GM145735 (CAW), NIH Diversity Supplement EY031710-01S1 (CAW); BBSRC APP26196 (DRO); Georgia Research Alliance (CRE). We thank Dr. Amir Vahabikashi at Northeastern University for scientific discussions and providing critical comments that greatly improved the quality of this work.

## Author Contributions

**Conceptualization:** S.M.S., W.D.S., D.R.O., C.R.E.; **Methodology**: S.M.S. (Designed the methods for pore formation assay), S.T.B. (Designed the methods for pore labeling assay); **Software**: S.M.S., B.M. (Designed the magnetic actuator using SOLIDWORKS); **Investigation**: S.M.S. (Investigated pore formation by seeding cells both atop and below micron-sized magnetic beads in phSC cell strains, HUVECs, and HDMVECs using widefield microscopy; Conducted confocal and scanning electron microscopy to detect pores; Investigated the effect of magnetic force on pore formation in phSC cells, HUVECs, and HDMVECs; Investigated the effect of Y-27632 treatments on pore formation in phSC cells, HUVECs, and HDMVECs while seeding cells below micron-sized magnetic beads; Investigated the effect of particle size and coating on pore formation), H.L. (Investigated pore formation in different phSC cell strains using widefield microscopy; Investigated the effect of Y-27632 and S1P treatments on cell stiffness and pore formation), B.M. (Investigated pore formation in HDMVECs; Built the magnetic actuator), C.A.W. (Measured cell stiffness using AFM), I.T. (Investigated bead-induction of pores), K.M.P. (Isolated and characterized phSC cells); **Validation**: S.M.S.; **Data Curation**: S.M.S.; **Formal Analysis**: S.M.S., H.L., B.N.S.; **Writing – Original Draft**: S.M.S., H.L., C.R.E.; **Writing – Review & Editing**: B.N.S. (Provided critical comments on magnetic force measurements and cell deformation), M.R.B.F. (Provided critical comments on surface treatment of particles and glass substrates), A.T.R. (Provided critical comments on SEM methodology), J.A.B., L.S. (Provided critical comments on cell culture methodology); **Critical Review**: W.D.S., D.R.O.; **Visualization**: S.M.S.; **Supervision**: S.M.S., C.R.E.; **Funding Acquisition**: W.D.S., D.R.O., C.R.E.; **Resources**: C.R.E., W.D.S.; All authors have reviewed, edited, and approved the final version of the manuscript.

## Supplementary Information

**Fig. S1.**
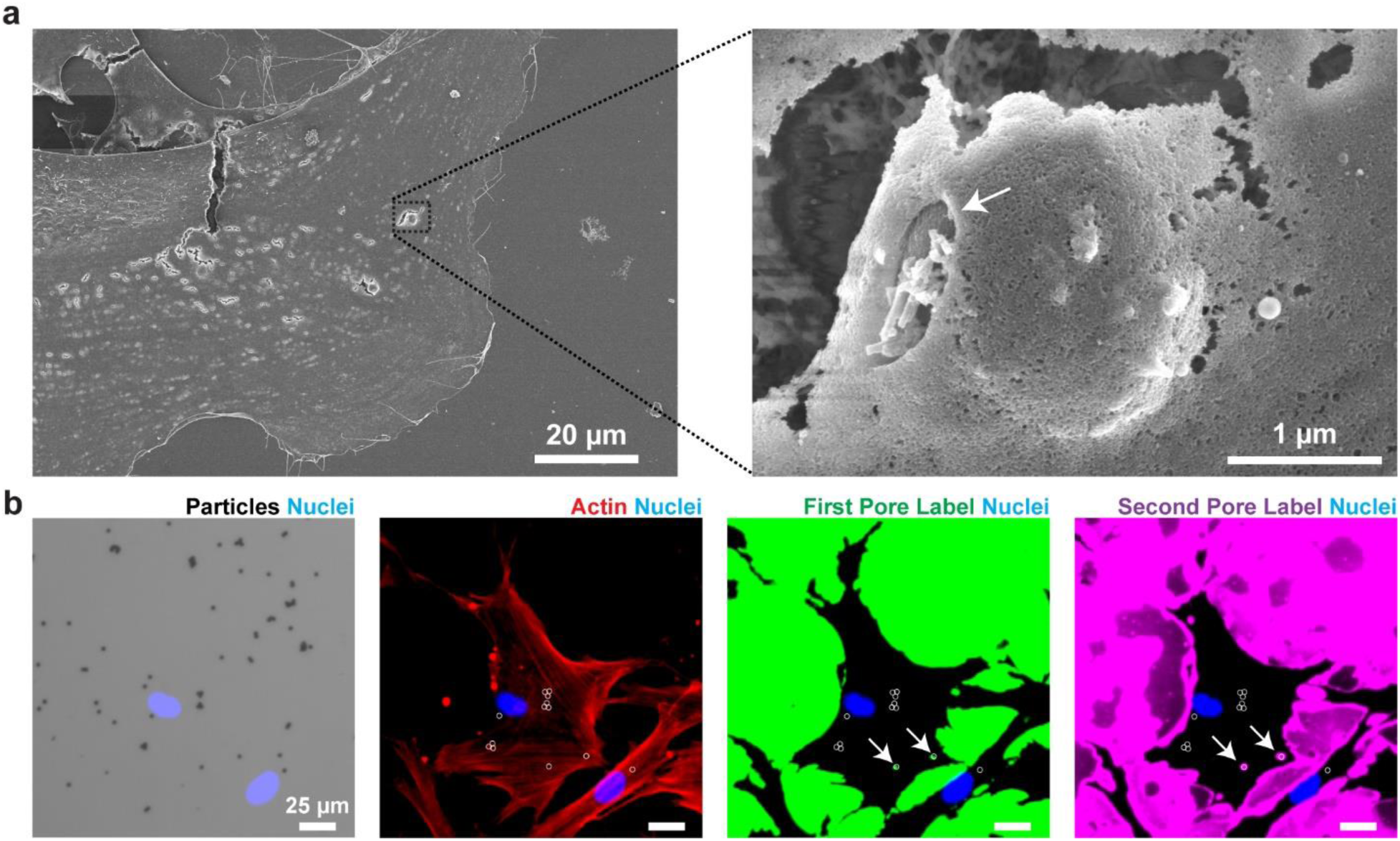
Transcellular pores induced by 2 µm particles seeded under phSC cells. **a,** SEM micrographs demonstrating particle-induced transcellular pore formation. The left image shows a low-magnification view of a phSC cell, while the right image shows a higher-magnification view of the inset (black box), highlighting the particle location and transcellular pore (white arrow). **b,** Brightfield and fluorescent images showing transcellular pores (white arrows) at the location of 2 µm particles. Pores were detected by the fluorescent assay (described in **Fig. 4a**) without any treatment between the first and second fluorescent tracers. The left image shows DAPI-stained nuclei (blue) merged with a brightfield image in which particles are seen as black dots. The particles below cells are shown by white circles in other images. Actin network (red), first fluorescent label (green), and second fluorescent label (magenta) are shown in second, third, and fourth columns, respectively. Arrows indicate locations of tracer accumulation coinciding with particles.

**Fig. S2.**
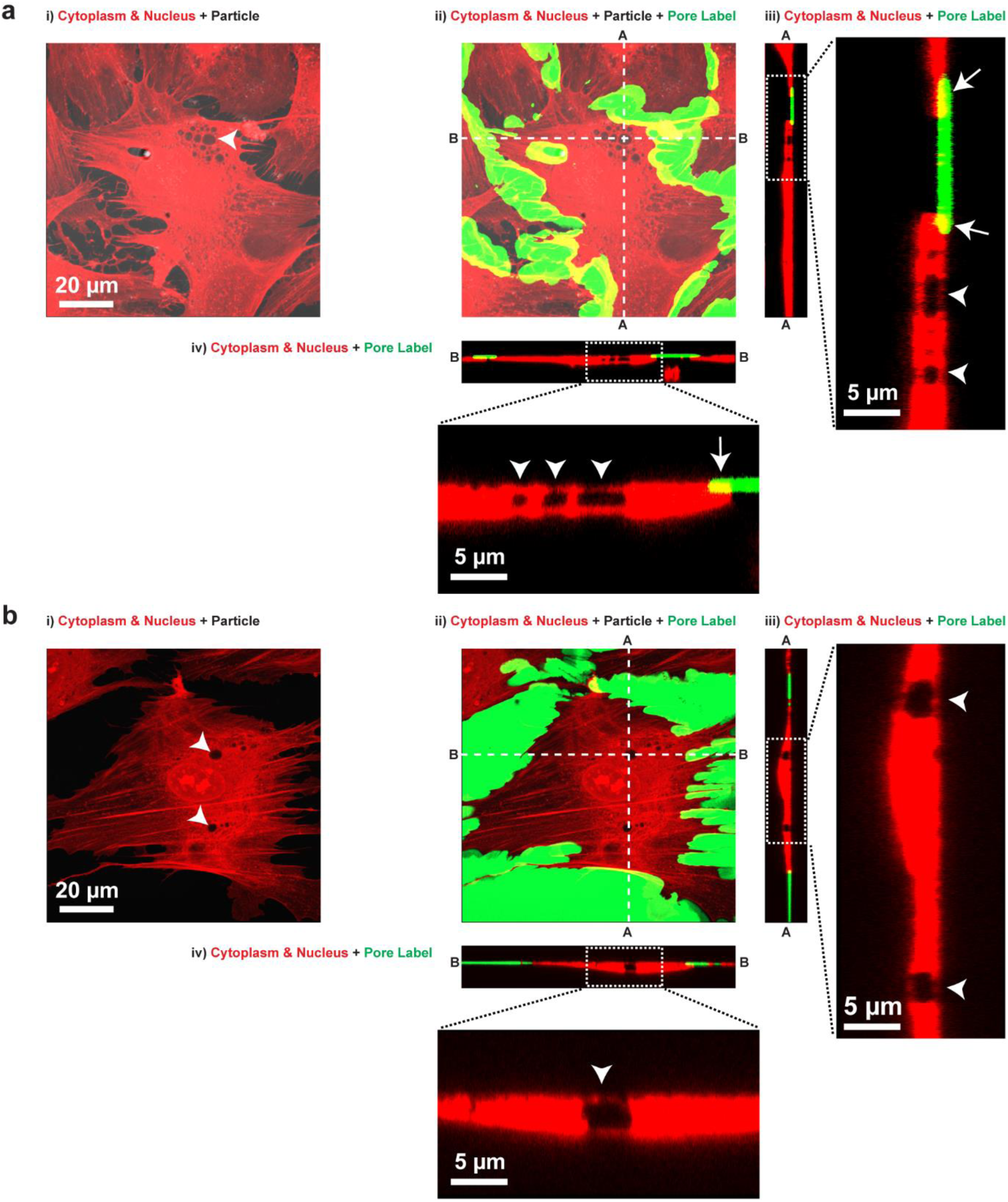
Intracellular vesicles detected in confocal images of cells seeded above a particle (a) and without any particles (b). In both **a** and **b**, subpanels **i** and **ii** show top views, maximum intensity projections of a phSC cell labeled for cell cytoplasm and nucleus (red), while subpanels **iii** and **iv** show cross-sectional side views of the cells at the location of intracellular vesicles (white arrowheads), along the A-A and B-B dashed lines in subpanels **ii**. Note that fluorescent tracer (shown in green in subpanels ii-iv) is bound to the biotinylated gelatin substrate outside of the cell footprint, but also slightly extended under the cell at the cell boundaries (white arrows). The cell shown in panel **a** is the same cell shown in **Fig. 3g**.

**Fig. S3.**
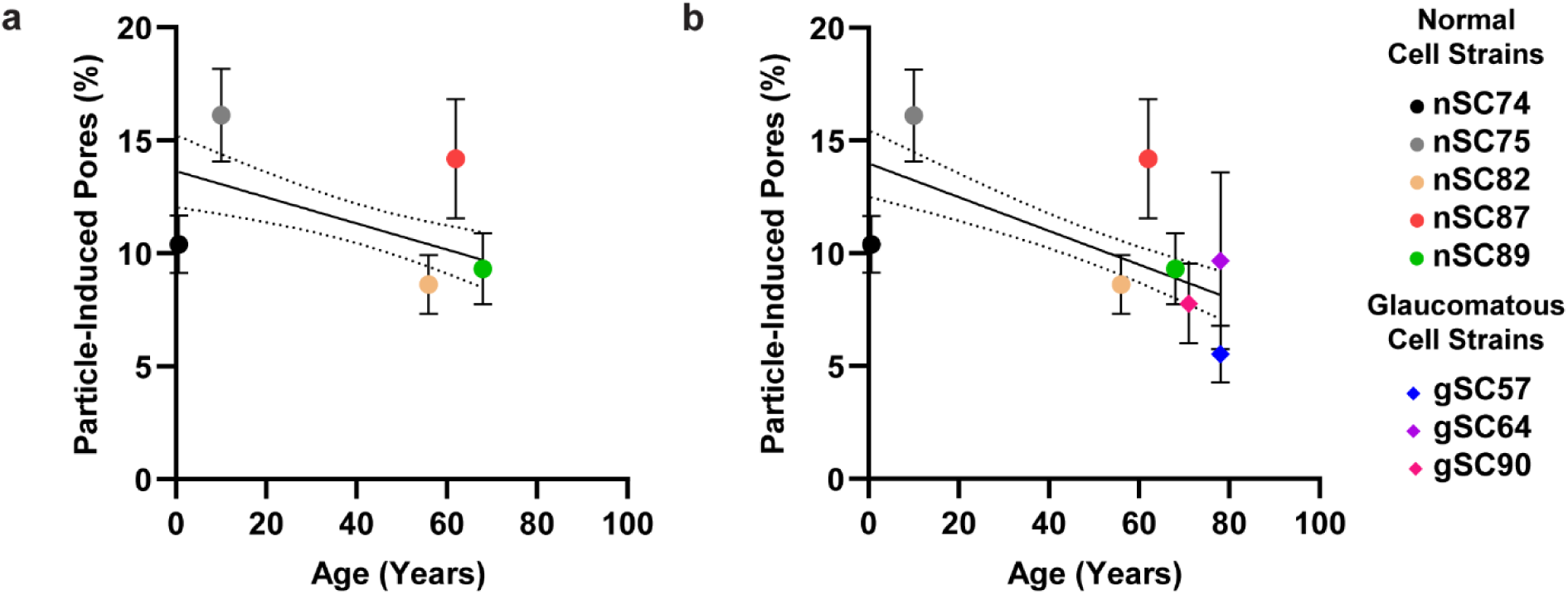
Weak association between pore formation and donor age in phSC cells. **a,** Simple linear regression demonstrated a weak relationship between particle-induced pore incidence and the age of normal phSC cell donors (R^2^ = 0.10, p = 0.0006). **b,** Three cell strains derived from older donors had a prior history of glaucoma. When accounting for glaucoma status as an independent variable, multiple linear regression revealed a somewhat stronger relationship between particle-induced pore incidence and the age of both normal and glaucomatous phSC cell donors (R^2^ = 0.39, p = 0.0004). Dotted lines represent 95% confidence intervals. Data are shown as mean ± standard deviation.

**Fig. S4.**
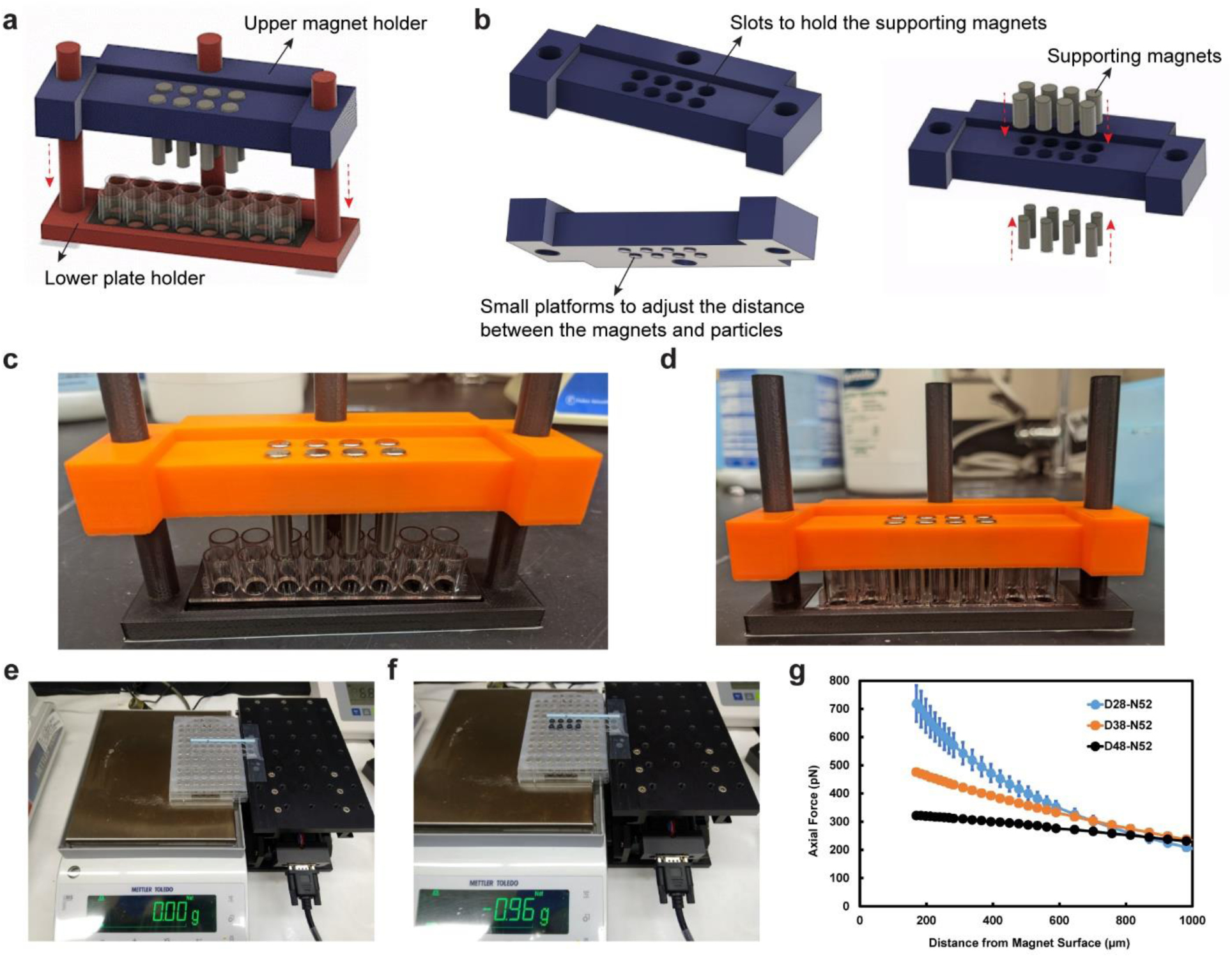
Design and functionality of custom magnetic actuator. **a,** A 3D computer-aided design illustrates the magnetic actuator, comprising a lower plate holder and an upper magnet holder. The upper magnet holder can slide down (indicated by red dashed arrows) to position the magnets at a controlled distance from the particles within the 16-well chambered slide. The device is engineered to exert magnetic force exclusively on the particles in the eight middle wells, while the remaining eight wells serve as control wells without magnets. **b,** Various views of the upper magnet holder are presented. Small spacers on the bottom surface of the upper magnet holder regulate the distance between the magnets (3/16” diameter x 1/2” thick, D38-N52, K&J Magnetics) and the particles, while eight slots on the top surface of the upper magnet holder accommodate supporting magnets (1/4” diameter x 1/2” thick, D48-N52, K&J Magnetics) securing the D38-N52 magnets in position. Final configurations are shown in **c** with magnets outside the wells and **d** with magnets inside the wells. **e**-**f,** The experimental setup for measuring the magnetic force exerted on particles is demonstrated. Magnets are placed in a well-plate on a scale and zeroed (**e**). Using a high precision Z-axis stage, a coverslip is positioned slightly above the magnets without direct contact. A known quantity of particles is then pipetted onto the coverslip above the magnets, and the scale is read (**f**) to calculate the axial force applied to the particles (see Methods). By incrementally adjusting the distance between magnets and particles using the Z-axis stage, the magnetic force applied on particles at different distances from the magnet surface is measured. **g,** Graph of average measured applied force on each particle versus the distance from the magnets. Not visible due to their small size are x-axis error bars of ± 20 µm, which was the step size of the positioning system. Three types of magnets were tested, with D28-N52 magnets (1/8” diameter x 1/2” thick, K&J Magnetics) exhibiting the highest force at shorter distances. However, due to their smaller diameter, the number of cells and particles in a well that were well-aligned with the magnet was limited. Consequently, only D38-N52 magnets were utilized in experiments requiring application of magnetic force.

**Fig. S5.**
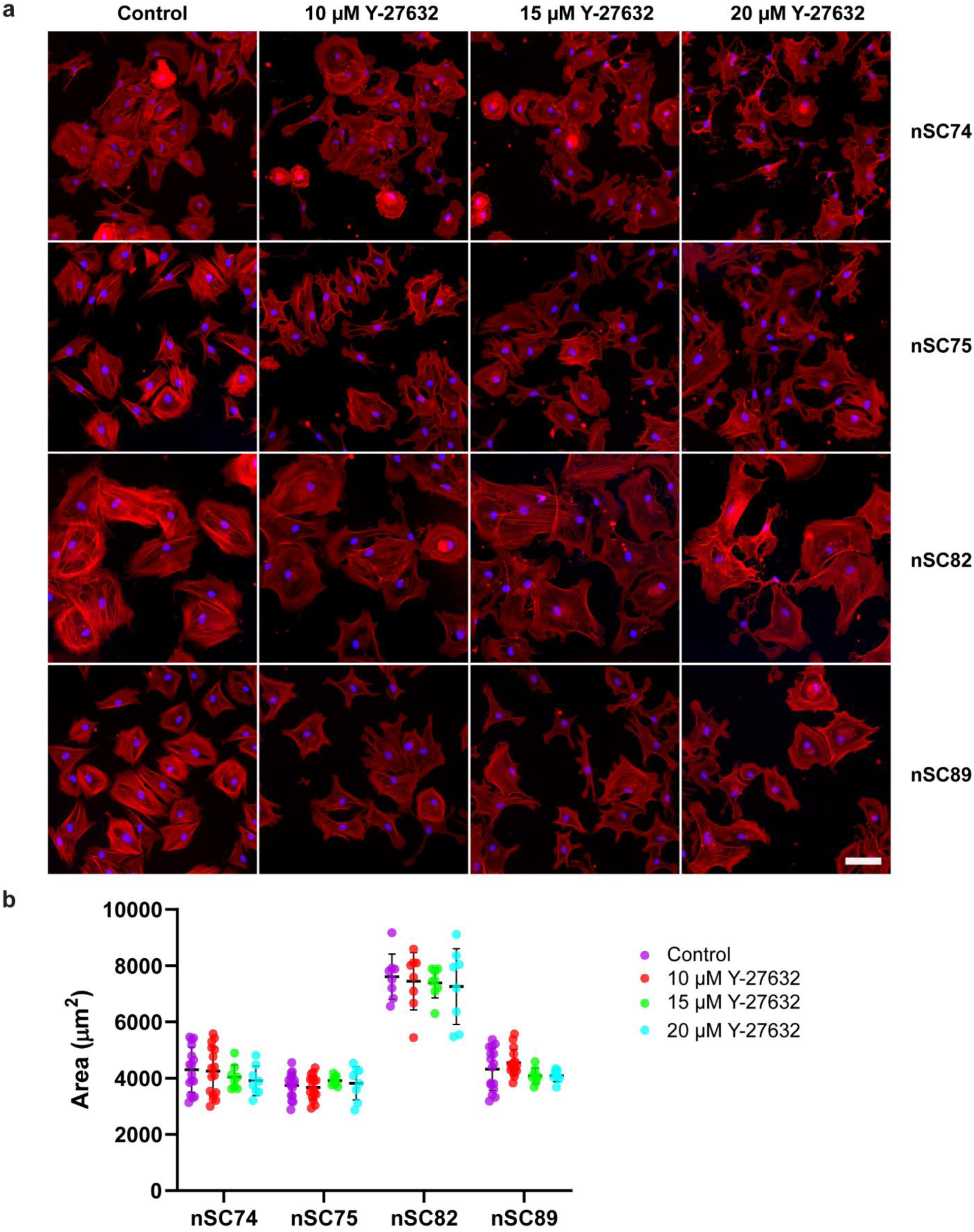
Impact of Y-27632 on cell area. **a,** Fluorescent images depict the actin network (red) and cell nuclei (blue) in nSC74, nSC75, nSC82, and nSC89 cell strains following Y-27632 treatments. Scale bar is 100 µm. **b,** Y-27632 treatments showed no significant effect on the area of cultured phSC cells. Analysis via 2-way ANOVA yielded the following P values: interaction = 0.8, cell strain factor < 0.0001, Y-27632 factor = 0.4.

**Fig. S6.**
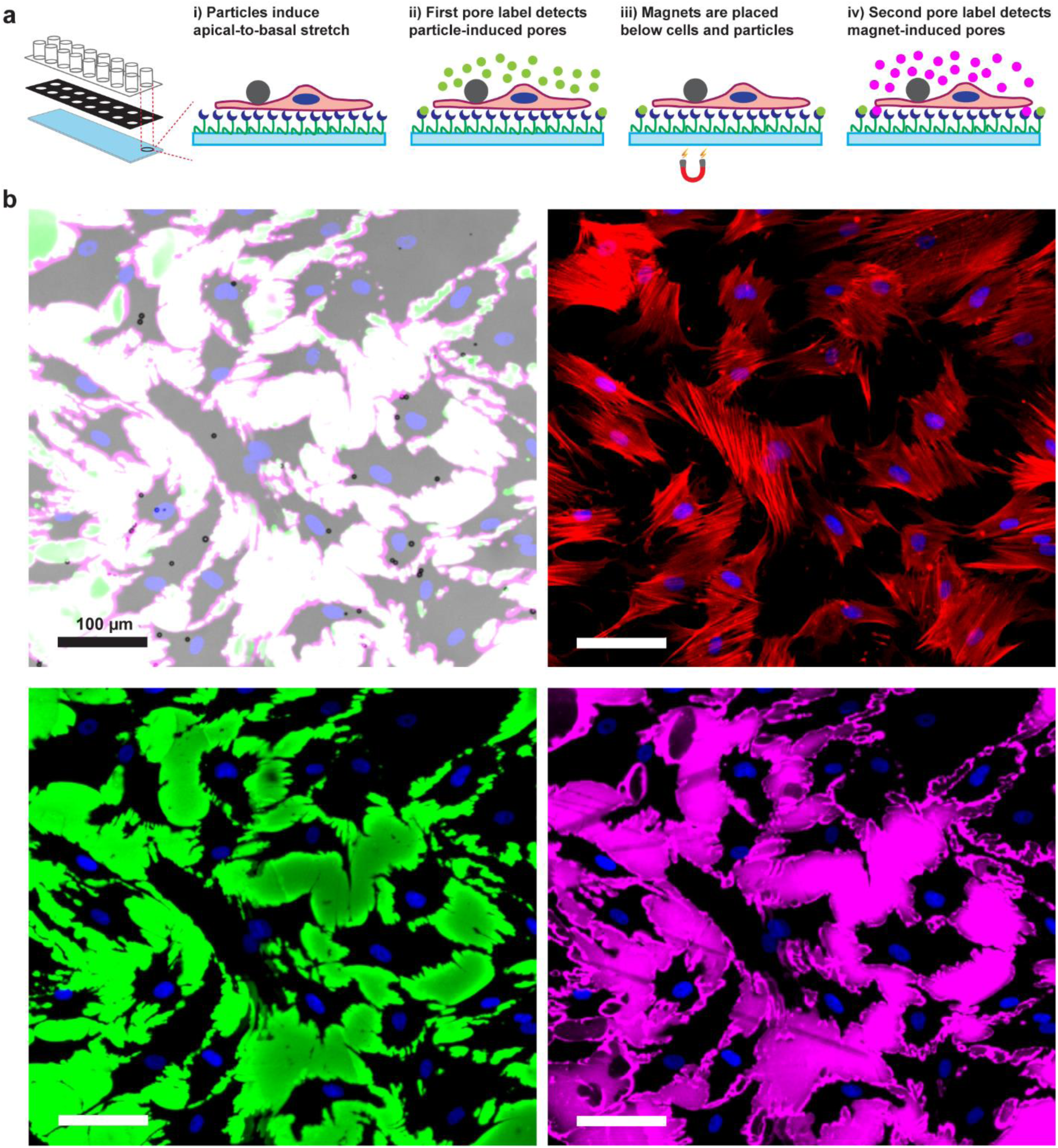
Transcellular pores were not detected when particles were placed on top of phSC cells. **a,** Illustration of the assay setup in which phSC cells were cultured over a biotinylated gelatin substrate in 16-well chambered slides, allowing for high-content detection of transcellular pores. Once cells were fully spread, particles were randomly seeded on top of the cells. A first fluorescent tracer (green) was applied to detect any transcellular pores formed solely due to particles. Subsequently, particles were subjected to a magnetic field to induce local apical-to-basal cellular deformation, thereby promoting pore formation. Magnet-induced pores at the particle locations were identified using a second fluorescent tracer (magenta). **b,** The top left image displays a brightfield image with particles represented as black dots, merged with fluorescent images of phSC cells exhibiting DAPI-stained nuclei (blue), the first fluorescent tracer (green), and the second fluorescent tracer (magenta). Other subpanels depict nuclei merged with the actin network (red), the first fluorescent tracer, or the second fluorescent tracer. No transcellular pores were detected at the particle locations. Treatment with 10 µM Y-27632 instead of applying a magnetic field also failed to induce transcellular pores.

**Fig. S7.**
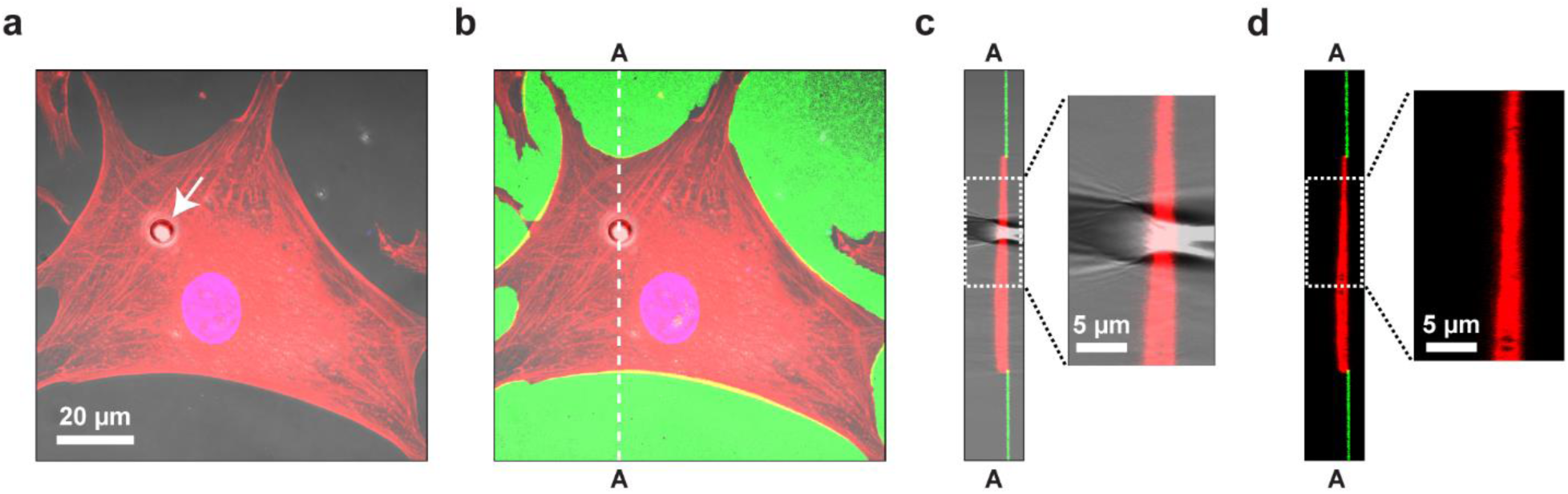
High resolution confocal images of particles placed on top of phSC cells. When particles were placed atop phSC cells, cell deformation was not observed, and transcellular pores were not detected. **a,** Top view, maximum intensity projections of a phSC cell labeled for cell cytoplasm (red) and nucleus (blue). The cell is spread on the biotinylated gelatin substrate while a particle (seen in the overlain bright field channel; white arrow) is positioned atop the cell. **b,** The fluorescent tracer (green) is bound to the biotinylated gelatin substrate outside of the cell footprint. **c,** Cross-sectional side view of the cell at the location of the particle (along the dashed line A-A in panel **b**), illustrating the particle’s position on top of the cell. Note that the optical distortion introduced by the particle disperses its image in the vertical direction. **d,** Similar to **c** but without the particle, highlighting the absence of cell deformation. The absence of fluorescent tracer at the particle location indicates that no transcellular pore was formed.

**Fig. S8.**
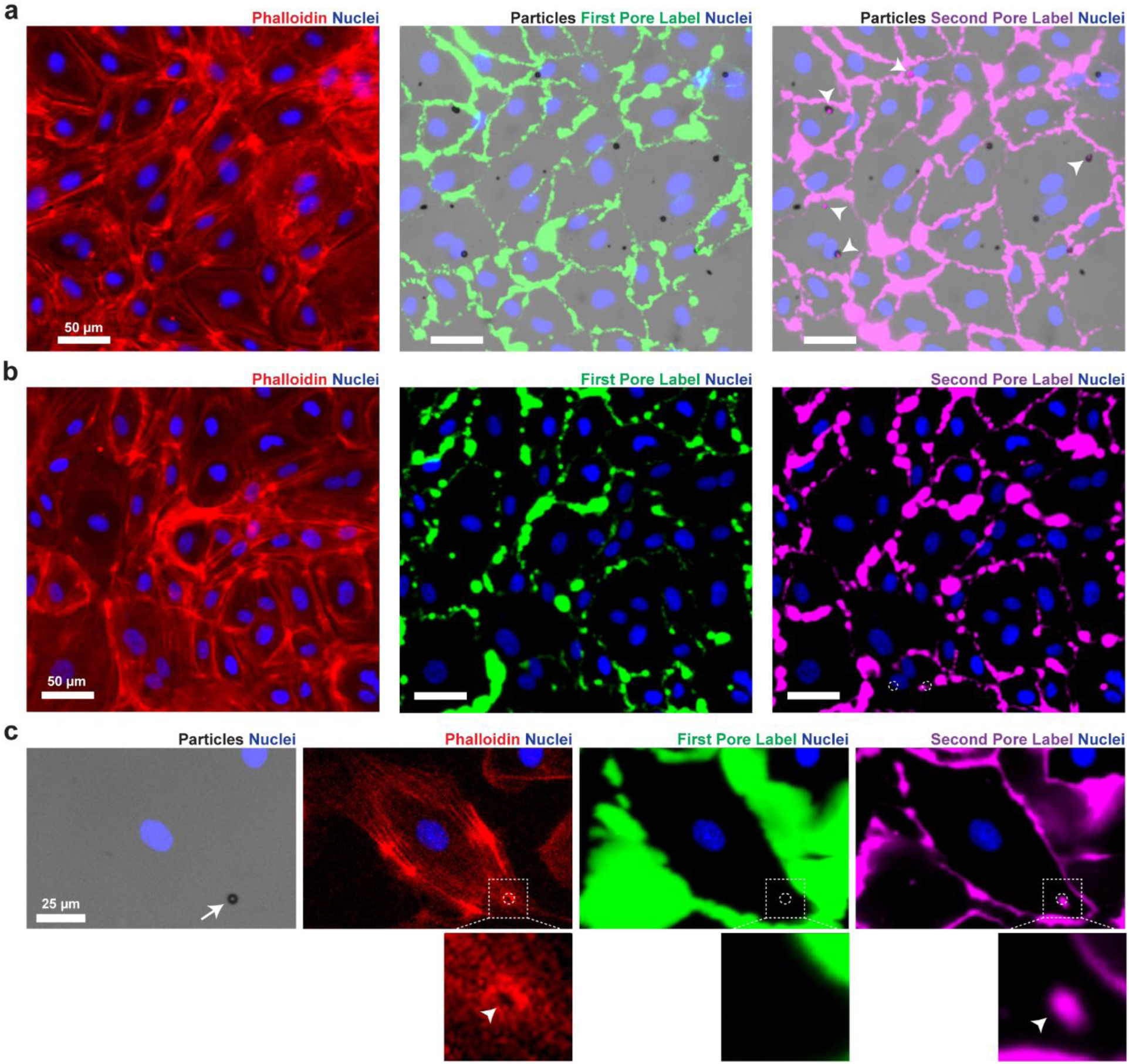
Induction of transcellular pores in human umbilical vein endothelial cells (HUVECs) and human dermal microvascular endothelial cells (HDMVECs) by magnetic particles placed on cell surfaces. HUVECs and HDMVECs were seeded with magnetic particles which were then exposed to a magnetic field, as depicted schematically in **Fig. S6a**. **a,** Low magnification brightfield and fluorescent images reveal transcellular pores in HUVECs where particles were positioned atop cells. Pores (white arrowheads) predominantly formed following the application of a magnetic field (473 ± 11 pN per particle). Please note that we did not quantify the number of pores before and after applying the magnetic field in these experiments. **b,** In control experiments, no transcellular pores were observed when no particles were added atop HUVECs. **c,** Higher magnification images depict a transcellular pore (white arrowheads) in HDMVECs. A particle (white arrow) is visualized as a black dot in the brightfield image. The location of the particle is also indicated by white circles in fluorescent channels of panel **c**. Insets in panel **c** provide magnified views at the location of the particle. Note the actin remodeling and the second fluorescent tracer at the location of particle. DAPI-stained nuclei are depicted in blue, the actin network in red, the first fluorescent tracer in green, and the second fluorescent tracer in magenta.

**Table. S1.** Total count of particle-induced (Fig. 5b), transient (Fig. 5d), emergent (Fig. 5f), and magnet-induced pores (Fig. 5g and h). This data is also presented in **Fig. 5** after normalization by the number of particles under cells.

|  | Particle-Induced Pores | Transient Pores | Emergent Pores | Magnet-Induced Pores |  |  |  |
| --- | --- | --- | --- | --- | --- | --- | --- |
|  |  |  |  | No Force | Low Force | Medium Force | High Force |
| Normal | 8607 | 2282 | 315 | 38 | 72 | 85 | 132 |
| Glaucoma | 1021 | 285 | 63 | 24 | 60 | 50 | 74 |

